# The landscape of immune dysregulation in pediatric sepsis at a single-cell resolution

**DOI:** 10.1101/2024.01.17.576030

**Authors:** Fahd Alhamdan, Sophia Koutsogiannaki, Koichi Yuki

**Author notes:** Correspondence: Koichi Yuki, M.D., Department of Anesthesiology, Critical Care and Pain Medicine, Cardiac Anesthesia Division, Boston Children’s Hospital, 300 Longwood Avenue, Boston, MA, 02115, USA.

## Abstract

Recognizing immune dysregulation as a hallmark of sepsis pathophysiology, leukocytes have attracted major attention of investigation. While adult and pediatric sepsis are clinically distinct, their immunological delineation remains limited. Breakthrough of single cell technologies facilitated the characterization of immune signatures. We tackled to delineate immunological profiles of pediatric sepsis at a single-cell level by analyzing blood samples from six septic children, at both acute and recovery phases, and four healthy children. 16 single-cell transcriptomic datasets (96,156 cells) were analyzed and compared to adult sepsis dataset. We showed a unique shift in neutrophil subpopulations and functions between acute and recovery phases, along with examining the regulatory role of resistin. Neutrophil signatures were comparable between adult and pediatric sepsis. Innate-like CD4 T cells were predominantly and uniquely observed in acute phase of pediatric sepsis. Our study provides a thorough and comprehensive understanding of immune dysregulation in pediatric sepsis.

## Introduction

Sepsis is the leading cause of death in intensive care unit (ICU) and the most expensive condition treated in the United States (U.S.).^1^ Over one-third of children who die in U.S. tertiary care pediatric ICU (PICU) have severe sepsis^2^. The Sepsis, Prevalence, Outcomes and Therapies (SPROUT) study, the first world-wide prospective study of pediatric sepsis, showed that the hospital morality of severe sepsis was significantly high (25%)^3^, comparable to adult sepsis. 67% of patients had multiple organ dysfunction syndrome (MODS) at the time of diagnosis, and 30% subsequently developed new or progressive MODS. Furthermore, recent reports suggest that its prevalence is on the rise^2,4,5^. These data hint the immediate need to decipher the underlying pathophysiology of pediatric sepsis for the improvement of our care.

Two important differences need to be noted between pediatric and adult sepsis. First, pediatric immune system is subjected to developmental changes^6^. Second, adult and pediatric sepsis differ in predisposing diseases and sites of infection^7,8,9^. However, current sepsis management is the same for both; primarily supportive, including fluid resuscitation, early antibiotic therapy, and cardiorespiratory support. Leukocytes are major players to fight against invading microorganisms. The contribution of immune dysregulation to morbidity and mortality has been described in adult sepsis^10^. However, there is a limited literature describing leukocyte phenotypic characteristics in pediatric sepsis. With the urgent need to develop sepsis therapeutics, delineating the immunological difference between pediatric and adult sepsis would allow us to develop specific treatment for each. Here we studied pediatric leukocyte transcriptomic signatures in pediatric sepsis using single-cell RNA sequencing (scRNA-Seq) and compared them to the ones from adult sepsis.

## Methods

### Human sepsis and control subject enrollment

In this study, we examined leukocyte signatures in control and sepsis pediatric subjects between May 2021 and January 2023. The study was approved by the Institutional Review Board at Boston Children’s Hospital, and written informed consent was obtained from all patients’ family. If indicated, assent was also obtained. The study was registered in ClinicalTrials.gov (Trial number NCT04103268) and carried out in accordance with Declaration of Helsinki. We targeted patients from 1 month of age to < 18 years old.

### Sample collection and leukocyte purification

As control subjects, we enrolled otherwise healthy patients who underwent elective procedures. In case of sepsis patients, we obtained blood samples at acute and recovery phases. We excluded patients with significant comorbidities such as known immune deficiency. 1 mL of blood was collected into heparin anticoagulant tubes. Leukocytes were purified by combining polymorphonuclear and mononuclear cell layers using Polymorphprep reagent (ProteoGenix, Schiltigheim, France) at room temperature. Leukocytes were immediately fixed with 10x fixing reagent (10x Genomics; Pleasanton, CA) and stored in liquid nitrogen until sequencing.

### 3’RNA library preparation and sequencing

Following a fast thaw in 37 °C water bath, single cell RNA libraries were generated using the Chromium Single Cell 3′ kit (10× Genomics). The cells were counted and loaded onto the Chromium Controller. Loading was performed to target capture of ∼3000 Gel beads-in-emulsion (GEMs) per sample for downstream analysis, and samples were processed through the Chromium Controller following the standard manufacturer’s specifications. The sequencing libraries were evaluated for quality on the Agilent TapeStation (Agilent Technologies, Santa Clara, CA), and quantified using Qubit 2.0 Fluorometer (Thermo Fisher Scientific, Waltham, MA). Pooled libraries were quantified using qPCR (Applied Biosystems, Waltham, MA) prior to loading onto an Illumina sequencing platform. The samples were sequenced at a configuration compatible with the recommended guidelines as outlines by 10× Genomics.

### Single-Cell RNA-Seq data analysis

Cell Ranger Software Suite (Version 3.1.0) (10× Genomics) was utilized to pre-process the raw sequencing reads. Following converting bcl files into fastq format, fastq files were de-multiplexed using mkfastq command. Subsequent UMI and cell barcode deconvolution along with mapping to the human reference genome GRCh38 were performed to generate final digital gene expression matrix files.

Single sample files with the h5 extension were further processed using Seurat package (Version 4.9.9.9045). Pre-filtering was applied on the Seurat objects with minimum 3 cells and 200 RNA transcripts. Subsequently, cellular mitochondrial content (MT) greater than 10% was filtered out from all Seurat objects. Following metadata build-up, all Seurat objects were merged and normalized. The rest of the processing steps including dimension reduction and unsupervised clustering were performed as in the default Seurat pipeline and by using 0.3 clustering resolution to exhibit major cell types.

Data integration and potential technical bias correction were computed with Harmony package (Version 0.1.1) according to donor, sequencing batch, gender and age parameters. Identification of differential expressed and cluster-specific marker genes was computed with FindMarkers and FindAllMarkers, respectively, implemented with Wilcoxon Rank Sum test. Marker genes utilized for all cell annotations were reported in the result section. For specific cell sub-clustering (neutrophils, monocytes, T cells, and B cells) a technique called community detection embedded in the Monocle 3 package (Version 2.26.0) was used for this purpose.

Trajectory analysis to probe neutrophils at differential states was performed by defining cluster 1 as root cells. Default settings were followed to generate gene modules in a resolution of 0.05. To infer autocrine and paracrine cell clusters’ interaction, CellChat package (Version 1.6.1) was used with default settings.

Functional analysis including biological pathways and processes were curated with the implementation of different databases such as Reactome, Biological process (Gene Ontology), NCATS BioPlanet^11–13^ by utilizing clusterProfiler package (Version 4.6.2). Cell cycle, phenotype, and function scores were computed with Seurat function cellcyclescoring and its dependency addmodulescore, the genes utilized in these analyses were imported from the literature or the above-mentioned pathway databases.

To predict potential surface protein markers from gene expression, SurfaceGenie shinyapp was used for the top markers of the cell cluster of interest (Neutrophils cluster 9)^14^.

### Phagocytosis

The neutrophil phagocytosis was done using Phagotest Kit (Glycotope Biotechnology; Heidelberg, Germany). Whole blood from donors were incubated in complete RPMI 1640 on ice for 10 min, followed by the addition of *FITC-E. Coli*. Then whole blood was kept on ice as cold control or was put into 37°C water bath for 10 min. At the end of incubation, samples were transferred back on ice, quenched, and washed. Neutrophils were stained with CD15-APC. Then, cells were subjected to lysis and fixation, and measured by BD Accuri C6 (BD Biosciences, Franklin Lakes, NJ).

### TLR4 reporter assay

Human TLR4 reporter cell line HEK-TLR4 cells containing a nuclear factor κ-light-chain-enhancer of activated B cells (NFκB)-inducible secreted embryonic alkaline phosphate (SEAP) reporter (InvivoGen; San Diego, CA) were used to assess the effect of resistin on TLR4 activation as previously reported^15^. The cells were cultured per the company’s protocol and stimulated with resistin (endotoxin filtered) at 0.1, 1 and 1 µg/mL (PeproTech, Cranbury, NJ) for 18 hours. The degree of NFκB activation was assessed by quantitating SEAP in the medium. Samples were subjected to a spectrophotometer analysis at 588 nm.

### TNF expression assessment in DMSO-differentiated HL60 cells and primary neutrophils

HL60 were differentiated into a neutrophil-like phenotype by incubating with 1.3% DMSO for 6 days as previously described^16^. The maturity of differentiated HL60 cell was probed by CD11b expression on flow cytometry using anti-CD11b-FITC antibody (BD Biosciences). Differentiated HL60 cells were harvested and stimulated with 10 µg/mL of resistin for 16 hours. Similalry, primary neutrophils were purified using magnetic beads and stimulated with resistin (10 µg/mL) for 6 hours. RNA was extracted with RNeasy plus mini kit (Qiagen, Hilden, Germany) according to the manufacturer’s instructions. Two-step cDNA synthesis kit was used to make cDNA. Real time quantitative polymerase chain reaction (RT-qPCR) was performed using SYBR Green (Applied Biosystems, San Francisco, CA) were used to assess TNF expression with the following forward and reverse primers (forward; CCTCTCTCTAATCAGCCCTCTG, reverse; GAGGACCTGGGAGTAGATGAG).

### Murine cecal ligation and puncture sepsis model

Mice were purchased from the Jackson Laboratory (Bar Harbor, ME, USA) and inbred in our animal facilities. All mice were on the C57BL/6 background and housed under specific pathogen-free conditions with 12-hour light/dark cycles. All the experimental procedures complied with institutional and federal guidelines regarding the use of animals in research and were approved by Boston Children’s Hospital Institutional Animal Care and Use Committee. Polymicrobial abdominal sepsis was induced by cecal ligation and puncture (CLP) surgery, as we previously performed^17^. In brief, 8-10-week-old wild type male mice were anesthetized with an intraperitoneal injection of ketamine 60 mg/kg and xylazine 5 mg/kg. Following its exteriorization, the cecum was ligated at a 1.0 cm from its tip and subjected to a single, through- and -through puncture using an 18-gauge needle. A small amount of fecal material was expelled with a gentle pressure to maintain the patency of puncture sites. The cecum was reinserted into the abdominal cavity. 0.1 mL/g of warmed saline was administered subcutaneously. Buprenorphine was given subcutaneously to alleviate postoperative surgical pain. Mice were euthanized at indicated time points for subsequent RNA sequencing experiments.

### Blood neutrophil isolation, RNA extraction and sequencing

Murine whole blood was centrifuged to obtain whitish buffy coat layer. This was then suspended in PBS and cells were stained with anti-mouse Ly6G, CD11b, and CD45.2 antibodies (Swamydas et al. Curr Protoc Immunol 2015; 110: 3.20.1-15). Neutrophils were flow cytometry-sorted as Ly6G^+^/CD11b^+^/CD45.2^+^ population. RNA was extracted from neutrophils using TRIzol Reagent (Thermo Fisher Scientific). RNA samples were quantified using Qubit 2 Fluorometer (Thermo Fisher Scientific) and RNA quality was assessed with Agilent TapeStation (Agilent Technologies). SMART-Seq v4 Ultra Low Input kit for Sequencing was used for full-length cDNA synthesis and amplification (Clonetech, Moutain View, CA), and Illumina Nextera XT library was used for sequencing library preparation, according to the manufacturer’s instructions. The final library was assessed with Qubit 2.0 Fluorometer and Agilent TapeStation. The sequencing libraries were multiplexed and clustered on one lane of a flowcell. After clustering, the flowcell was loaded on the Illumina HiSeq instrument according to the manufacturer’s instructions.

### Bulk RNA-Seq data analysis

Data processing was performed with Galaxy platform. Quality of sequencing reads was assessed by FastQC v0.72. Reads were further aligned to the mouse reference genome (Mus Musculus mm10) with RNA STAR v2.7.8a. Mapped reads were counted with featureCounts v2.0.1, followed by exon DESeq2 v2.11.40.6 analysis to determine differentially expressed genes. Z-scores in heatmaps and pairwise time-point comparison utilized the variance stabilizing transformation (VST) normalized gene expression generated with DESeq2 tool. Bulk RNA-Seq deconvolution analysis was conducted using CIBERSORTx^18^. A signature matrix was retrieved from our neutrophils datasets after converting the human genes to their corresponding murine orthologs, mixture files were constructed from the VST normalized counts of the CLP samples at the two time points.

### Statistical analysis

Statistical analysis was performed using GraphPad Prism 9 software package (GraphPad Software Inc., San Diego, CA). The type of statistical methods used was described under each figure legend. P< 0.05 was considered statistically significant.

## Results

### Neutrophilia and lymphopenia as two key phenomena of pediatric sepsis

Our clinical cohort included six children with sepsis and four healthy children (**Fig. 1a**). Sepsis samples were collected at both acute and recovery phases. Demographics and pediatric Sequential Organ Failure Assessment (pSOFA) scores of the clinical cohort were depicted in **Suppl Fig. 1a.**

**Fig. 1:**
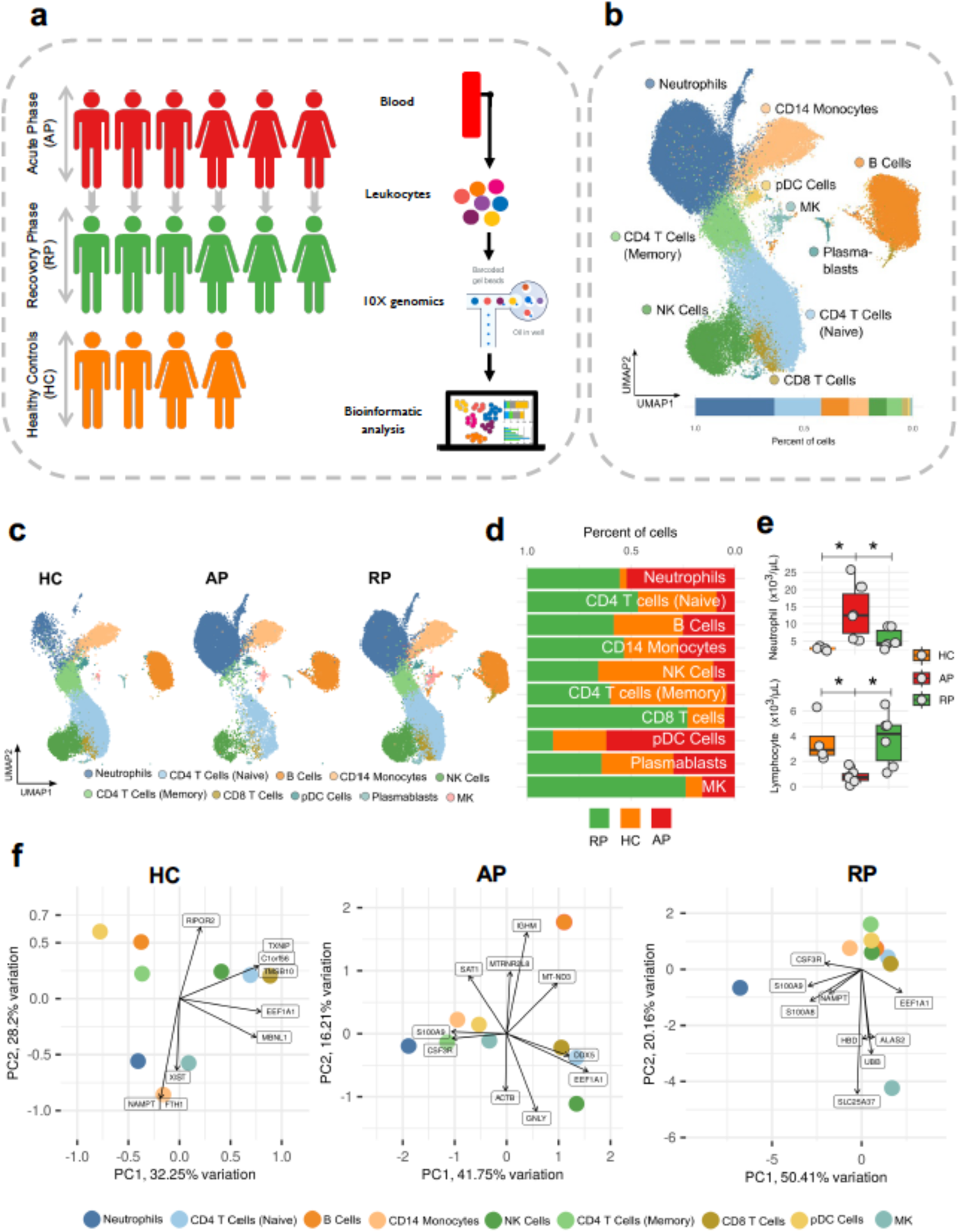
Major cell types of circulating leukocytes in pediatric sepsis. **a,** Schematic representation of the study design. Acute phase: AP (*n* = 6), recovery phase: RP (*n* = 6) and healthy controls: HC (*n* = 4). **b,** Uniform Manifold Approximation and Projection (UMAP) for the major circulating leukocyte cell types with their corresponding annotations and partitioning. **c,** UMAPs and bar plots depicting the abundance differences across the three conditions of study subjects. **d,** Boxplots showing the comparison of the lymphocyte and neutrophil counts among the three study conditions. Data were shown as mean ± S.D. One-way ANOVA with Bonferroni *post hoc* analysis. * p< 0.05. **e,** Principal Component Analysis (PCA) of the top 500 variable genes among all the cell types with the significant gene loadings.

To capture specific transcriptomic changes of pediatric sepsis at a single-cell resolution, we isolated peripheral blood leukocytes and performed single-cell sequencing on the Chromium platform. The 16 single cell datasets generated a total of 96,156 cells that were grouped into 12 cell clusters (**Suppl Fig. 2a**). We then annotated the cell clusters using different algorithms such as Azimuth^19^ and CellMarker^20^. Consequently, 10 major cell types were determined (**Fig. 1b**), showing their top three genes (**Suppl Fig. 2b**). We then asked if the major cell types exhibited any fractional abundance among the three subject conditions (healthy/steady, acute and recovery sepsis states). At a first glance, there was a drastic elevation of neutrophil fractions in acute phase (AP) and recovery phase (RP) compared to healthy controls (HC) (**Fig. 1c-e** and **Suppl Fig. 2c**). We also observed a remarkable reduction in the fractions of CD4^+^, CD8^+^ T cells, and NK cells in AP compared to the other two conditions. Cellular compositions for each pediatric subject, per age group and gender are illustrated in **Suppl Figs. 1b** and **2d-e**.

Neutrophil-to-lymphocyte ratio (NLR) has been frequently used as a prognostic tool for the outcomes of various diseases including adult sepsis ^21,22^. In pediatric sepsis, NLR at admission is suggested as a marker of severity with the cutoff of ∼4, in which high NLR score may indicate the severity of stress response^23^. In our clinical cohort all pediatric subjects demonstrated significantly elevated NLR scores in AP (22.6 ± 15.3) compared to RP (2.6 ± 3.0) or control (0.97 ± 0.51) (**Suppl Fig. 2f**), supporting the severity of our sepsis cohort.

To infer the dynamic changes in gene expression of the major cell types, we computed principal component analysis (PCA) using the top 500 variable genes in the three conditions. The PCA exhibited a distinct expression profile of neutrophils in RP, marked with *S100A8*, *S100A9*, *NAMPT*, and *CSF3R* (**Fig. 1f**). This distinct expression profile of neutrophils in RP was further verified by means of unsupervised hierarchal clustering (**Suppl Fig. 3a**). Of note, *NAMPT* is located at the downstream of granulocyte colony-stimulating factor (*G-CSF*) signaling, which facilitates the growth of new neutrophils^24^.

Because we observed a close proximity in the expression profiles of both neutrophils and CD4^+^ memory T cells in AP, we next determined to identify the phenotype of the CD4^+^ memory T cells. By excluding the cell-doublets probability (**Suppl Fig. 3b**), these cells represented an innate-like CD4^+^ T cell phenotype^25^, marked with *CSF3R* expression (**Suppl Fig. 3c**). Interestingly, the fractional abundance of the innate-like CD4^+^ T cell fractions decreased in older children (≥ 6 years) (**Suppl Fig. 3d**).

### Distinct neutrophils subtypes in pediatric sepsis phases

As aforementioned, neutrophils expanded in number in both sepsis phases. Therefore, we intended to gain a granular view of their gene expression profiles and subpopulations. To this end, we first conducted neutrophils sub-clustering that revealed new nine neutrophil subpopulations (**Fig. 2a**) with their associated top 10 genes (**Suppl Fig. 4a**).

**Fig. 2:**
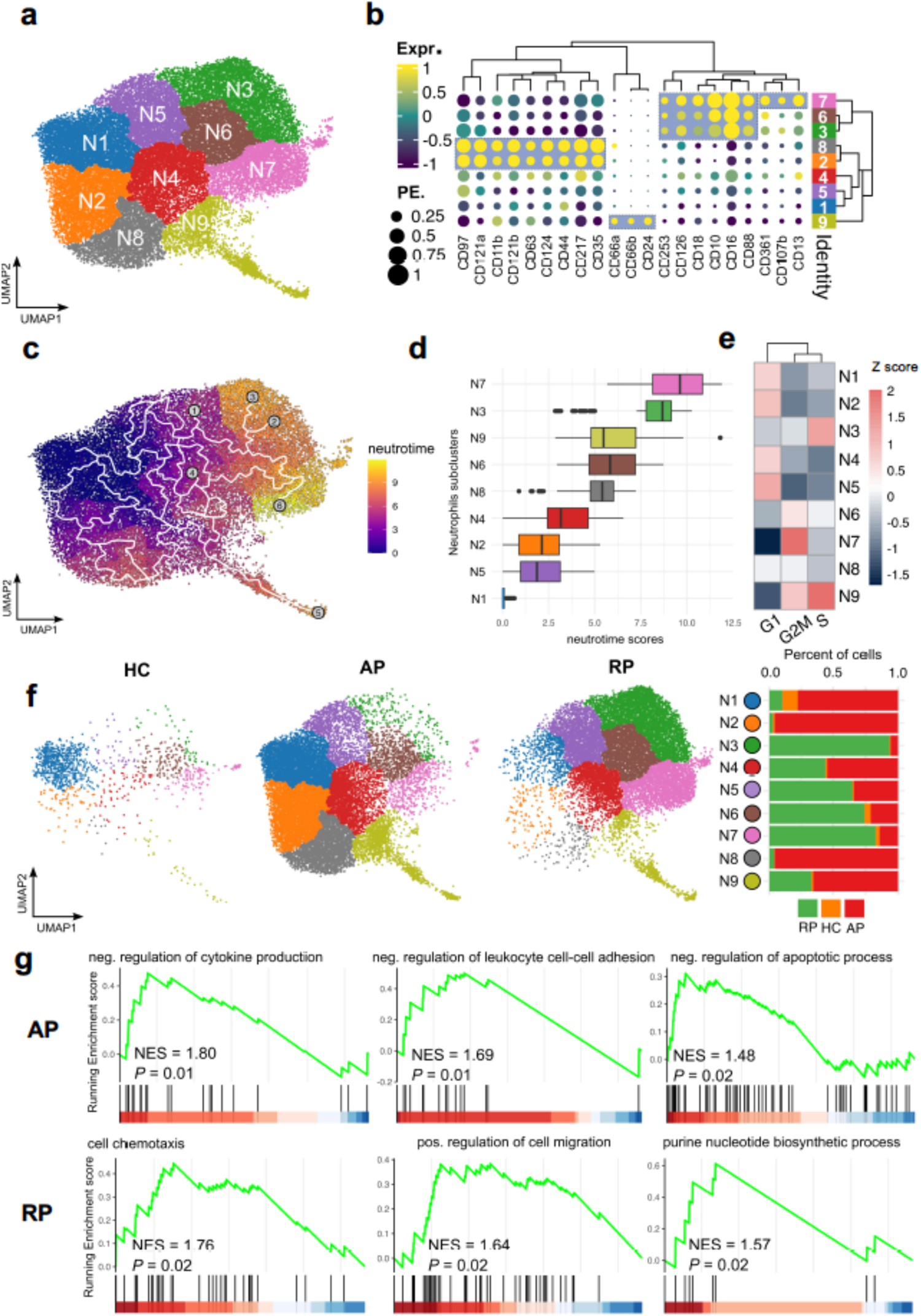
Neutrophil subpopulations with the trajectory analysis. **a,** UMAP of neutrophils subclusters for all 16 study individuals. **b,** Clustered dot plot of the expression of neutrophil marker genes, showing the aggregation of the neutrophil subclusters based on the expression of these markers. **c-d,** neutrotime trajectory analysis (c) of neutrophils from N1 to N7 with the corresponding neutrotime values in subfigure (d). **e,** Cell cycle scores (G1, G2M, and S) for each of the neutrophil subclusters represented as Z scores. **f,** UMAPs and bar plots for the three study conditions depicting the differences of abundance in neutrophil subclusters. **g,** Gene Set Enrichment Analysis (GSEA) displaying three significant Gene Ontology: Biological Processes (GO: BP) of differentially expressed genes between AP and RP.

To further characterize the nine subclusters, we curated a panel of surface CD-proteins based on their significant gene expressions for each subcluster (**Fig. 2b**). Six neutrophil subclusters were then aggregated into three subtypes, namely N2, N8 as (*IL1R2*) CD121b^+^ immature neutrophils, N9 as CD66b^+^ degranulating neutrophils and N3, N6 and N7 as CD16^+^ CD10^+^ mature neutrophils, indicating distinct differential fates of these neutrophil subclusters.

To further address this observation, we performed trajectory analysis through generating pseudo/neutro-time scores to define fate probability for each single neutrophil subcluster (**Fig. 2c-d**). Whereas the immature N2 and N8 were broadly overrepresented in the low neutrotime score group, the mature N3, N6 and N7 showed the highest differential capability together with the degranulating N9, belonging to a higher neutrotime score group. To support this outcome, cell-cycle dynamics demonstrated that N3, N6, N7 and N9 underwent active transcription, whereas N1, N2, N4 and N5 exerted low self-renewal ability (**Fig. 2e**).

We next inspected the fractional distribution of the nine neutrophil subclusters across the three subject conditions (**Fig. 2f**). Surprisingly, AP and RP revealed a balanced partitioning of N1, N2 and N8 (highly prevalent in AP) against N3, N6, and N7 (dominant in RP), respectively, albeit with drastic underlined biological differences. N4, N5, and N9 exhibited an overlapping pattern, displaying a shared cellular signature that bridged the boundary between AP and RP.

To gain a biological snapshot of neutrophils in AP and RP, we performed differential gene expression analysis of neutrophils in AP versus RP and inferred the top three biological processes. On the contrary to the active and migratory RP neutrophils (**Fig. 2g**), AP neutrophils revealed negative regulation of cytokine production, leukocyte cell-cell adhesion and apoptotic process. In addition, we examined the phagocytic function in AP versus RP in *ex vivo* settings. A higher phagocytosis capacity was observed in RP (**Suppl Fig. 5**). These and the previous findings reflect an immunosuppressive status in AP of pediatric sepsis.

### Functional analysis of neutrophils subpopulations and role of resistin in pediatric sepsis

Next, we intended to inspect several biological functions and processes for each neutrophil subcluster. To this end, we utilized AddModuleScore algorithm to compute gene scores using several gene sets (Fig. 3a and supplementary table 1). As anticipated, N2 and N8 (neutrophil subclusters abundant in AP) showed lower maturation levels compared to N3, N6, and N7 (neutrophil subclusters abundant in RP). Although these neutrophils subclusters exhibited comparable activation levels, N2 and N8 deemed to be dysfunctional since their maturation, proliferation and migration scores were lower than in N3, N6 and N7. Additionally, essential neutrophils defense mechanisms such as phagocytosis, neutrophil extracellular traps (NETs) and reactive oxygen species (ROS) formation attained highest scores in N3 and N7 (**Fig. 3a**). In line with the trajectory analysis, aging score suggested N7 as the terminally differentiated form of neutrophils (**Fig. 3a**). Consistent with this finding, phagocytosis was prominent in RP together with protein phosphorylation and cell adhesion, while defense response and oxidative stress were attenuated in AP (**Suppl Fig. 4b**). This characterized the reversal of the immunosuppressive status in neutrophils between the two sepsis phases, thereby implying an ongoing cellular transition from AP to RP.

**Fig. 3:**
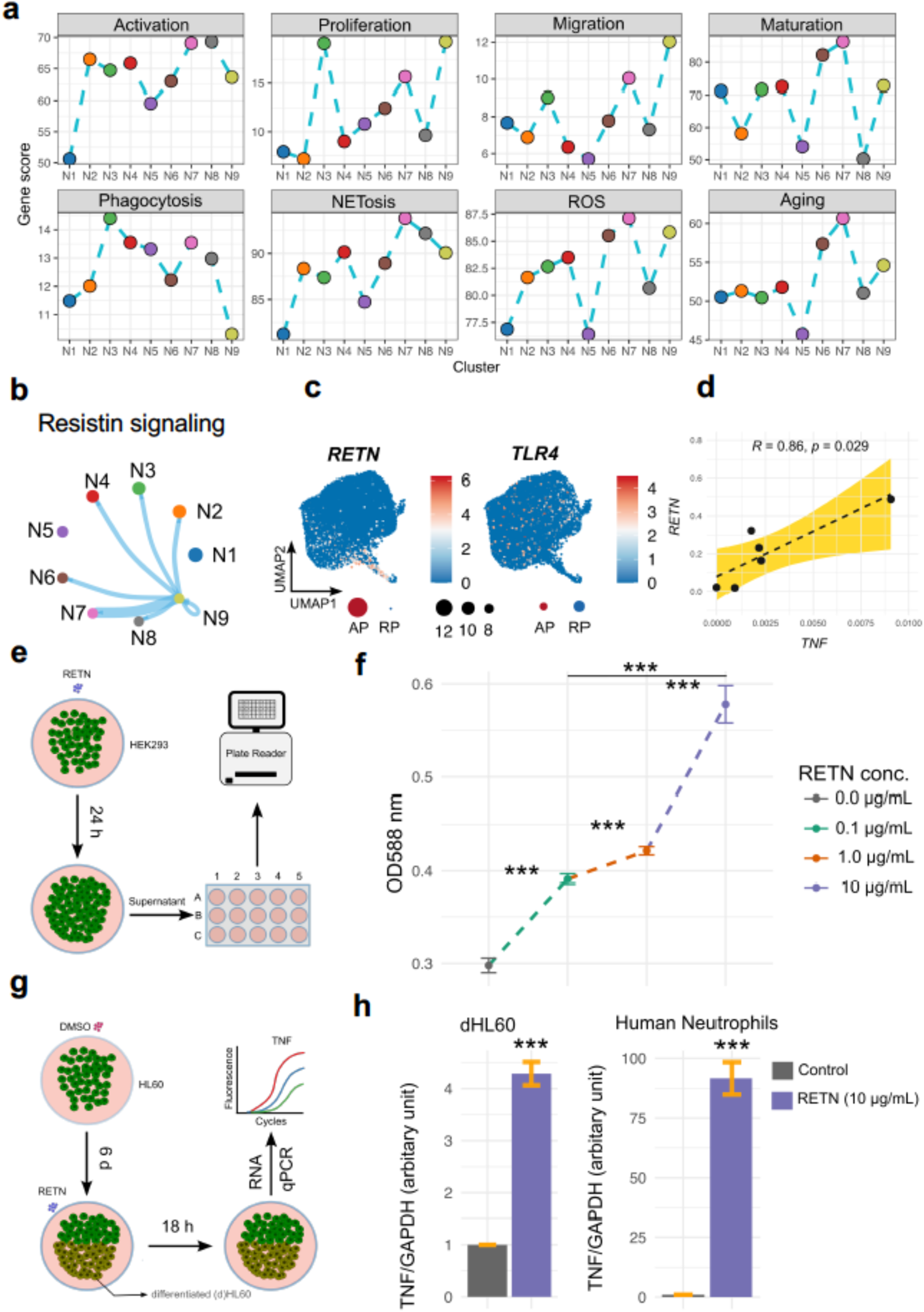
Functional analysis of the neutrophil subpopulations. **a,** Comparisons of functions and biological processes gene scores amongst the neutrophil subclusters computed via using a gene-set for each function or biological process illustrated in Supplementary Table 1. **b,** Cell-cell communication analysis showing the resistin signaling interaction between N9 and N9, N2, N3, N4, N6, N7. **c,** Feature and dot plots displaying the expression of *RETN* and *TLR4* genes. **d,** Linear regression analysis between the expression of *RETN* and TNF in all neutrophils for each subject. **e,** Schematic representation of the experimental setup for TLR4 reporter assay by treating HEK293-TLR4 cells with resistin. **f,** Line plot showing the fluorescence intensity scores in the HEK293-TLR4 cells supernatant following 24-hour incubation with resistin. Data were shown as mean ± S.D. One-way ANOVA with Bonferroni *post hoc* analysis was used for statistical significance. *** p< 0.001. **g,** Schematic representation of the experimental setup for HL60 cells by differentiating them into neutrophils using 1.3% DMSO and further treating them with resistin. **h,** Bar plot illustrating the mRNA expression of TNF in differentiated HL60 cells and human neutrophils stimulated with 10 µg/mL resistin. Data were shown as mean ± S.D. One-way ANOVA with Bonferroni *post hoc* analysis. *** p< 0.05.

To examine the cellular communication between the neutrophils subclusters, we carried out cell-cell interaction analysis based on receptor-ligand binding probability, which revealed that resistin (RETN) signaling interconnected subcluster N9 with most of neutrophil subclusters except for N1 and N5 (**Fig. 3b**). RETN presumably binds to the extracellular receptor TLR4^26^. The cell-cell interaction matrix matched the expression pattern of both *RETN* and *TLR4* (**Fig. 3c**), in which both were more elevated in AP.

To examine the role of RETN in TLR4 signaling, we first employed the TLR4 reporter assay *in vitro*. RETN was capable to activate TLR4 signaling in a dose-dependent manner (**Fig. 3e-f**). To examine the inflammatory tendency of RETN, we performed linear regression analysis, which showed a robust positive correlation between *RETN* and tumor necrosis factor (*TNF*) in AP (**Fig. 3d**). To test this *in vitro*, we differentiated HL60 cells into neutrophil-like phenotype via 1.3% DMSO stimulation (**Fig. 3g-h**)^16^. The differentiated (d)HL60 cells were further stimulated with RETN. Consistent with the reporter assay and linear regression, RETN significantly induced the expression of TNF in dHL60 cells, which was further validated in human neutrophils (**Fig. 3h**).

### Degranulating N9 as regulatory neutrophils in pediatric sepsis

To gain further insights into the functional role of the *RETN*-expressing N9 subcluster, we first identified the differentially expressed genes (DEGs) of RP versus AP. The majority of DEGs in N9 were enriched in RP including the N9 top nine genes (**Fig. 4a and Suppl Fig. 4a**), among which were *MMP8* and *LTF* representing the specific granules content. By performing granules enrichment analysis for N9, we identified that the degranulating phenotype was predominantly linked to RP and prominently represented by specific granules (**Fig. 4b)**, matching the moderate maturation level of N9. These findings were in agreement with the overall dysfunctional state of neutrophils during AP.

**Fig. 4:**
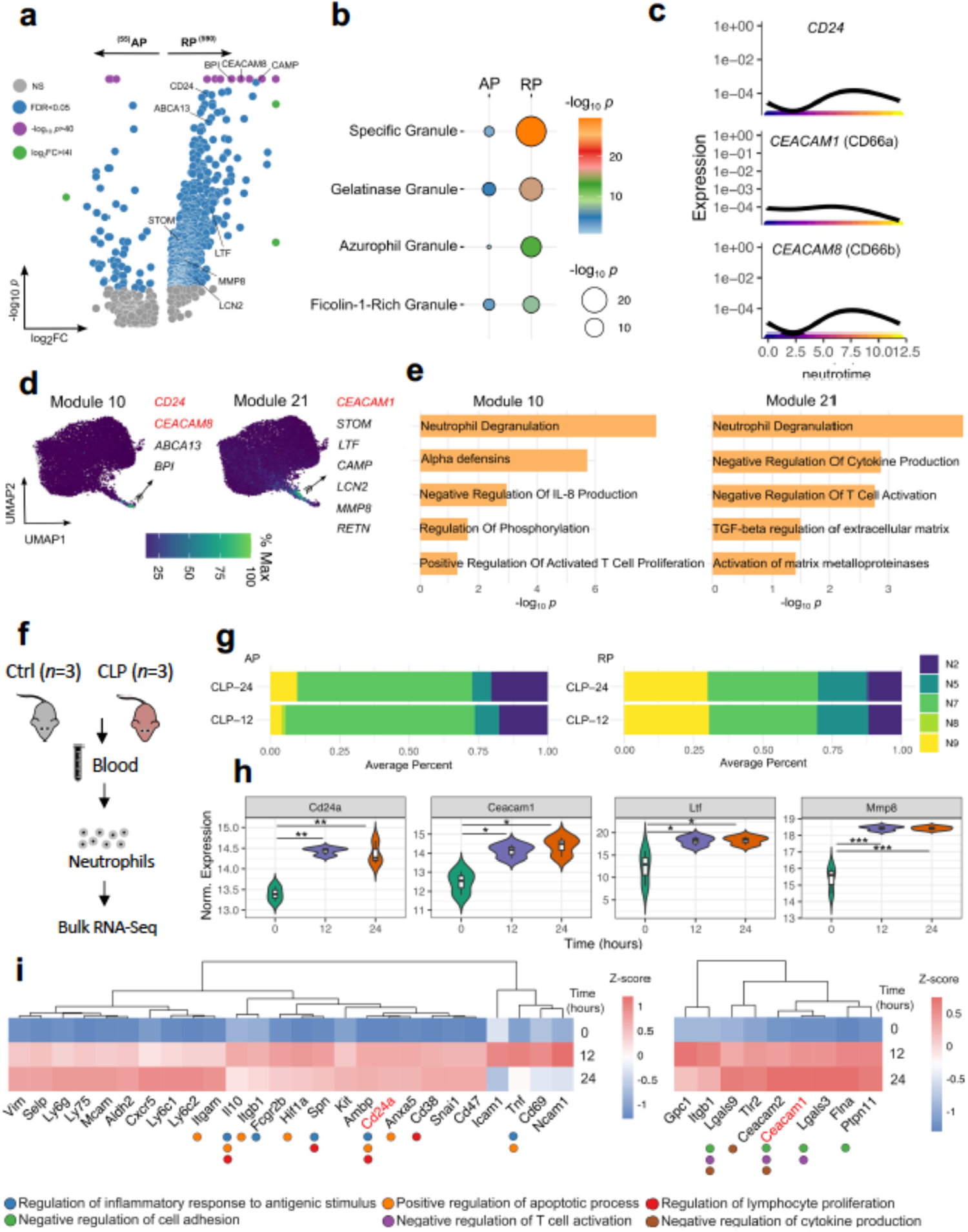
Characterization of the neutrophil subcluster N9. **a,** Volcano plot depicting the differentially expressed genes (DEGs) between AP and RP in N9 and by labeling the top nine genes. **b,** Dot plot for the enrichment analysis of four neutrophil granules significantly enriched in Gene Ontology-Cellular Compartment database. **c,** Expression dynamics of *CD24*, *CEACAM1* and *CEACAM8* along the neutrotime. **d,** Two specific gene modules of N9 neutrophils expressing the top three surface marker gene (*CD24*, *CEACAM1* and *CEACAM8*), co-expressing also the top ten 10 genes in N9. **e,** Significant biological processes and pathways associated with the genes expressed in the two N9 gene modules. **f,** Schematic representation of the experimental setup for the CLP murine model and bult RNA-Seq analysis of blood neutrophils. **g,** Bulk RNA-Seq deconvolution analysis depicting the average percentage of neutrophils subclusters in the CLP murine model. **h,** Box plots depicting the expression levels of Cd24a, Ceacam1, Ltf, and Mmp8. One-way ANOVA with Bonferroni *post hoc* analysis was used for statistical significance. \**P*□<□0.05; \*\**P*□<□0.005; \*\*\**P*□<□0.001. **i,** Heatmaps of the gene network of either Cd24a or Ceacam1 with their associated biological processes. Mouse genes were written in small characters.

The dynamic expression of the three N9 surface marker genes along the neutrotime demonstrated that *CEACAM1* (CD66a) was activated in an early stage during sepsis development (**Fig. 4b)**. This was previously observed in the surface markers panel as N8 shared the overexpression of this gene with N9 (**Fig. 2b**), indicating that CD66a represented a mixed-phenotype of neutrophils.

Next, we aimed to explore the biological functions of N9. Thus, we inspected co-varying genes among neutrophils and found 24 gene modules (**Suppl Fig. 6b**), among which two modules specifically enriched in N9 and contained the three surface marker genes *CD24*, *CEACAM1* (CD66a), and *CEACAM8* (CD66b) as well as the other N9 top ten genes (**Fig. 4c**). Consensus pathway analysis of the two gene modules revealed that neutrophil degranulation pathway was overrepresented in both analyses (**Fig. 4d**). Alpha defensins including *DEFA1B*, *DEFA1B*, *DEFA3*, and *DEFA4* were significantly enriched in gene module 10. Additionally, we observed a variety of canonical immune regulation pathways was associated with the genes of the two modules. These data indicated that N9 encompassed a paradigm of immune regulatory and defense mechanisms.

We next sought to validate the existence of the degranulating N9 subcluster in an experimental abdominal murine sepsis model induced by CLP (**Fig. 4f**). Following bulk RNA-Sequencing of whole-blood neutrophils, deconvolution analysis demonstrated a greater cellular enrichment of N9 in an RP phenotype at comparable abundances at 12 and 24 hours. Surface marker genes of N9 including Cd24 and Ceacam1 were significantly overexpressed at 12 and 24 hours after CLP surgery (**Fig. 4g**). Their overexpression was synchronized with the upregulation of the specific granule genes *Mmp8* and *Ltf* genes. This suggested the presence of the N9 neutrophil subcluster in the murine sepsis model.

To simulate the single-cell setup, we inferred the significant genes in our bulk RNA-Seq analysis that are known to interact with either Cd24a or Ceacam1 using the STRING database (**Fig. 4h**), by assuming that these genes belong to the same neutrophil subpopulation. Akin to the human data, a variety of immune regulation pathways was associated with the *Cd24a* and *Ceacam1* interacting genes in murine blood neutrophils. Taken together, this suggests that *CD24^+^ CEACAM1/8^+^* neutrophils might contribute to the immune regulation machinery in sepsis pathogenesis, in line with the concept of the N9 regulatory role.

### A massive immune dysregulation in T and NK cells in AP

In light of our earlier observations, the second phenotypic change in AP was lymphopenia (**Fig. 1d**). To further explore this, we subclustered T and NK cells into new cell subpopulations. 11 subclusters were reconstituted and annotated to their corresponding cell types (**Suppl Fig. 7a-c**). Following partitioning cells according to sepsis conditions (**Fig. 5a**), we observed a pronounced global cell depletion in AP, which was significantly aggravated in (CD8) T8, (CD4) T2, T4, T6, and T7 cells.

**Fig. 5:**
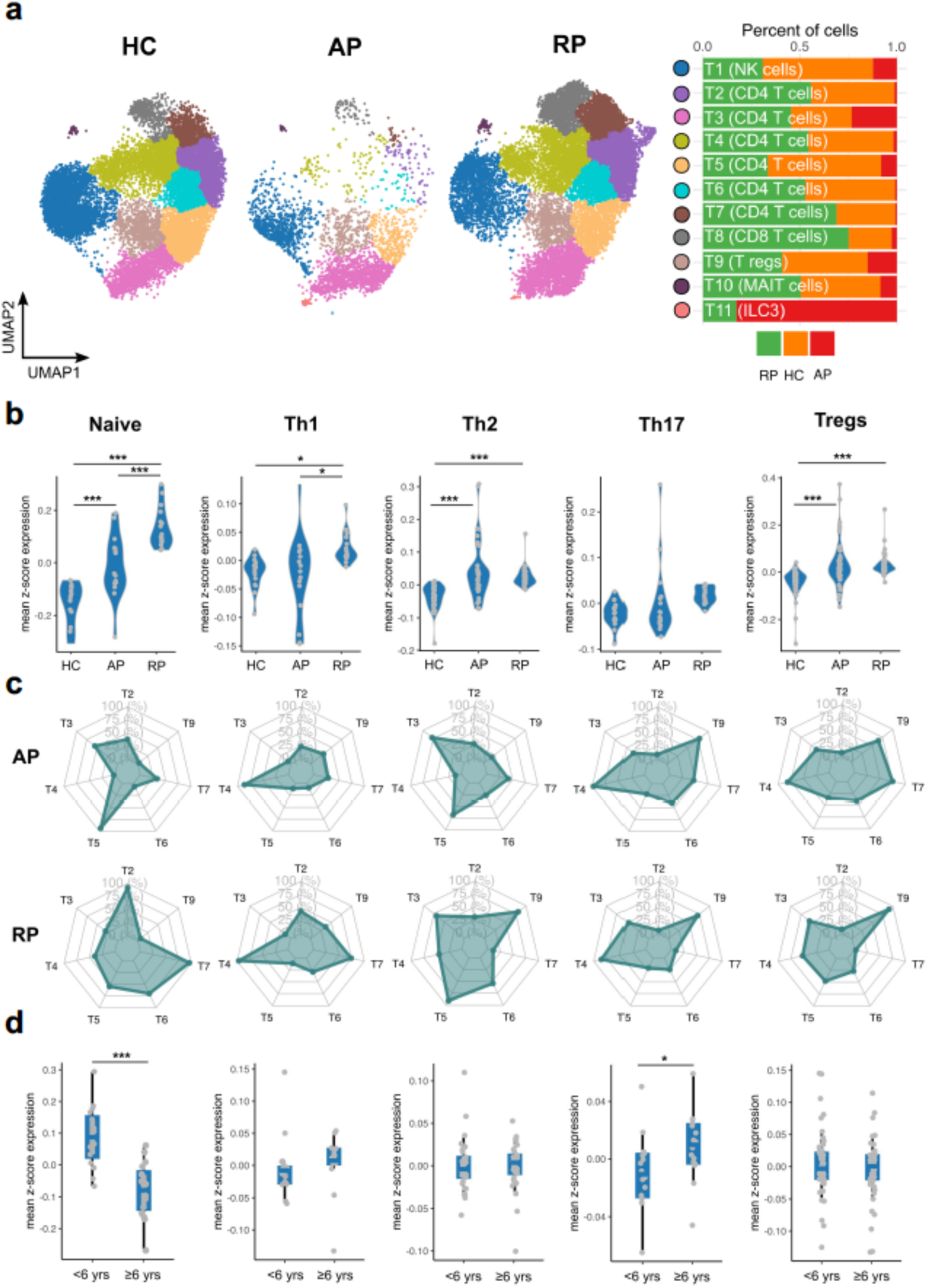
Severe depletion of T and NK cells in sepsis acute phase. **a,** UMAPs and bar plots for the three study conditions depicting the abundance differences in T and NK subclusters. **b,** Violin plots comparing gene signatures of Naïve, Th1, Th2, Th17 and Tregs between HC, AP, and RP. One-way ANOVA with Bonferroni *post hoc* analysis was used for statistical significance. \**P*□<□0.05; \*\**P*□<□0.005; \*\*\**P*□<□0.001. **c,** Radar plots showing the dynamic changes in the previous polarized T cell subsets across the subclusters of the CD4 T cells in AP and RP. **d,** Box plots comparing gene signatures of Naïve, Th1, Th2, Th17 and Tregs the two age groups. Unpaired student t test was used for statistical analysis. \**P*□<□0.05; \*\*\**P*□<□0.001.

To better understand the phenotypic diversity in CD4 T cell subpopulations, we inferred the gene scores of their polarized cell subsets in each sepsis phase and steady state (**Fig. 5b**). Interestingly, the naïve T cells were exponentially increasing from HC to RP through AP, perhaps in part due to the recruitment of a significant large naïve T cell pool to the infection site during sepsis ^27^. We additionally observed a steady expansion in regulatory T cells in AP and RP. Unexpectedly, Th17 polarized subset did not represent the predominant phenotype in the pediatric sepsis infection, but rather Th2 in both AP and RP synergized with Th1 in RP. The contribution of the single CD4 T cells subclusters to the Th-subsets were illustrated in **Fig. 5c**.

We then asked if these changes in the CD4 T cells polarization patterns were age-related. While Th1, Th2, and Tregs kept consistent phenotypic profiles, Th17 subset significantly restored its typical infection-related phenotype in older children (≥ 6 years) (**Fig. 5c**).

Given the robust alteration in T cell proportions and states, we assessed their versatile biological functions and processes (**Fig. 6a and Suppl Table 2**). As T3 and T5 represented a naïve T cell subset in AP and their activation frequencies were at lowest (**Fig. 5c)**. In contrast, T4 exhibited the highest activation, proliferation, differentiation and migration profiles.

**Fig. 6:**
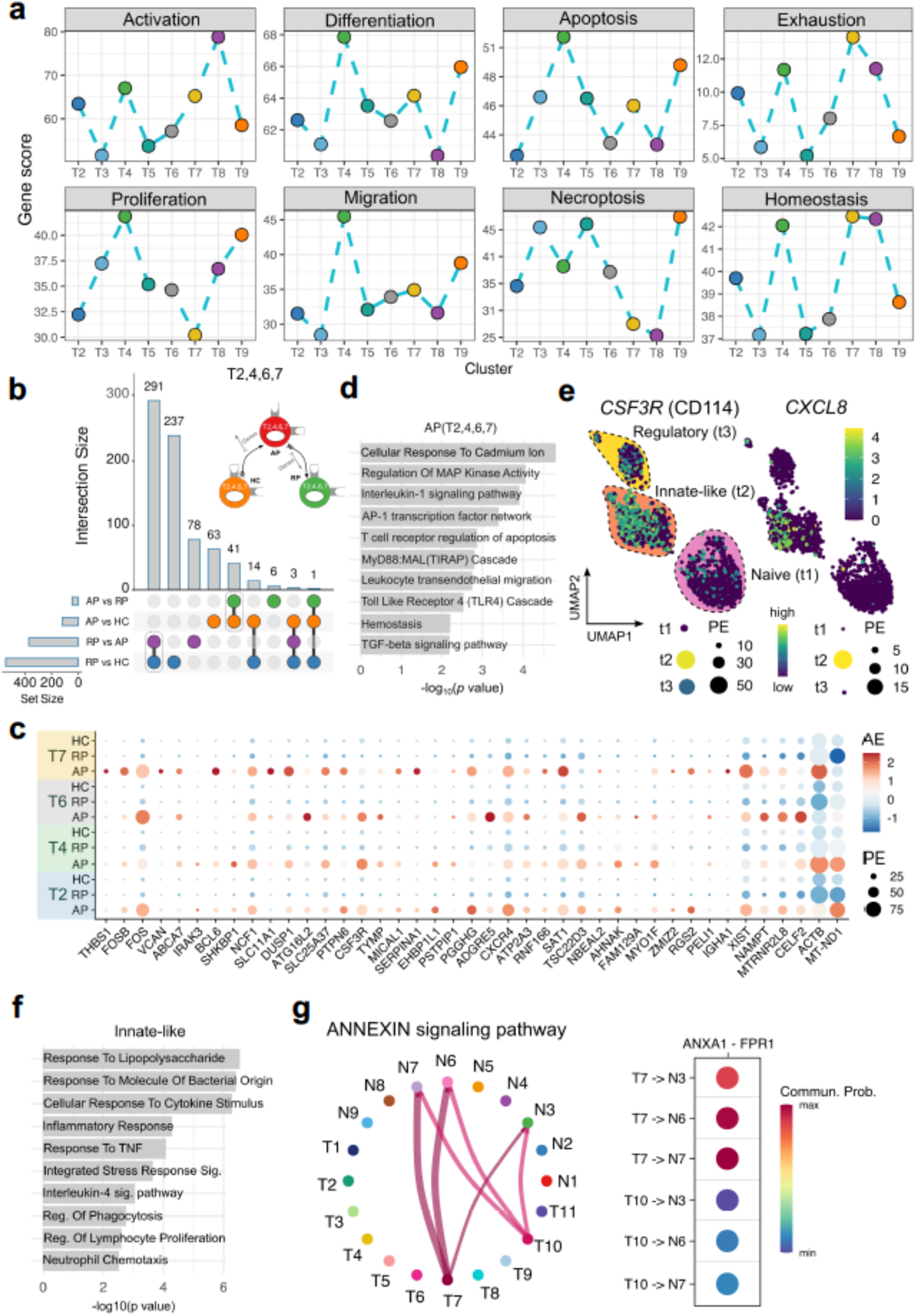
Innate-like T cells as significant signature for AP in the memory T cells compartment. **a,** Comparisons of functions and biological processes scores amongst the T cell subclusters computed via using a gene-set for each function or biological process illustrated in supplementary table 2. **b,** Upset diagram of the unique and shared DEGs (FDR□<□0.05) between a series of pairwise comparisons of HC, AP, and RP in severely-depleted CD4 T cell subclusters (T2, T4, T6, T7). **c,** Dot plot for the 41 DEGs that are upregulated and shared for AP vs RP and AP vs HC. **d,** Bar plot showing the significant biological processes and pathways associated with previous 41 DEGs. **e,** Feature plots for the expression of *CSF3R* and *CXCL8* as gene markers innate-like CD4 T cells. **f,** Bar plot showing the significant biological processes and pathways associated with the DEGs upregulated in the innate-like CD4 T cells compared with naïve and regulatory cell portions in AP. **g,** Cell-cell communication analysis showing the ANNEXIN signaling interaction between T7, T10, N3, N6, and N7 (right) and communication probability scores of the interaction between the previous subclusters (left).

Necroptosis, but not apoptosis notably culminated in T3, T5, and T9. These subclusters were depleted only modestly in AP. In contrast, severely depleted T2, T4, T6, T7 and T8 exhibited the highest exhaustion and homeostasis scores. T cell homeostasis is a restoring process to compensate for numeric loss of T cells following cell depletion^28^.

To investigate the pathways accompanied the depletion of CD4 T cells, we separately analyzed the DEGs of the severely depleted T2, T4, T6, T7 (**Fig. 6b-s**). 41 DEGs were uniquely upregulated in T2, T4, T6, T7 in AP compared with RP and HC (**Fig. 6c**). These DEGs and their associated biological processes and pathways predominantly represented an innate immune phenotype. Thus, we overlaid the cellular composition of CD4 T cells in AP (**Fig. 6e**). As anticipated, the innate-like CD4 T cells persisted within the memory CD4 T cell compartment despite its severe cell depletion in AP compared to RP and HC (**Fig. 6e** and **Suppl Fig. 7d**).

The innate-like CD4 T cells were first described by Gibbons et al as *CXC8L*-expressing CD4 T cells in newborns ^29^. In line with this, we observed *CXC8L* expression in this population (**Fig. 6e**). However, there was still a caveat in characterizing them based on their surface markers since some researchers described them as CD31^+^ T cells^29–31^, which was a blunt surface marker as evidenced by our data compared to *CSF3R* (CD114) expression levels (**Fig. 6e** and **Suppl Fig. 7e**). The biological processes of the innate-like CD4 T cells were associated with neutrophil chemotaxis, response to lipopolysaccharide (LPS) and TNF, and inflammatory response contrasted with typical biological processes for the other CD4 T cell fractions in AP (**Fig. 6f and Suppl Fig. 7e**). Consisted with the biological role of innate-like CD4 T cells, cell-cell interaction analysis revealed that T7 interconnected with the functionally mature N3, N6 and N7 neutrophil subclusters through Annexin signaling pathway. The latter promotes chemotaxis in neutrophils upon binding ANXA1 to the leukocytes chemotactic receptor FPR1^32^ (**Fig. 6g**).

### Lack of antigen processing and presentation by monocytes and B cells

The initial analysis of major cell types revealed a comparable cell fraction of monocytes and B cells between HC and AP (**Fig. 1c** and **Suppl Fig. 9a**). First, we first performed fine-tune clustering on monocytes and found three distinct subpopulations (**Suppl Fig. 8a**). Following a refined annotation of these subpopulations into classical, intermediate, and no-classical monocytes (**Suppl Fig. 9b**), we observed severe depletion of the intermediate monocytes in AP and a global absence of the nonclassical monocytes in HC (**Suppl Fig. 8a**).

Likewise, we subclustered B cells into five annotated subpopulations (**Suppl Fig. 8a** and **Fig. 9c**). CD27^+^ B memory cells experienced severe depletion in AP, while transitional B cells were totally vanished (**Suppl Fig. 8a**). Plasma and classical memory B cells were notably expanded in AP and RP compared to HC.

Following rigorous analysis of the DEGs across the subclusters of monocytes and B cells, we detected a common denominator between the depleted cells in the intermediate monocytes (M2) and transitional B cells (B2) in AP. These two subpopulations function as antigen presenting cells (APCs) mainly for CD4^+^ T cells as they overexpress HLA-DR family genes together with other antigen processing genes (**Suppl Fig. 8b**). This observation is considered as the third phenotypic change in the acute phase of our pediatric sepsis cohort.

Based on the previous observation, we sought to untangle additional cellular interactions among monocytes, T and B cells in AP and RP (**Suppl Fig. 8c**). Consequently, RETN signaling pathway was overrepresented in AP interconnecting monocytes with B and T subclusters. Contrary to AP, CD4^+^ T and mucosal associated invariant T (MAIT) cell compartments in RP were identified to be the source of Annexin and macrophage migration inhibitory factor (MIF) signaling pathways that further regulated various monocytes, T and B subpopulations.

### Comparison of adult and pediatric sepsis

To the best of our knowledge, this study was the first to investigate pediatric sepsis at a single-cell resolution. Therefore, we conducted a comparative analysis between pediatric and adult sepsis. To this end, we reanalyzed publicly available single-cell data of peripheral blood mononuclear cells (PBMCs) from two healthy donors and five sepsis adult patients (3 survivors and 2 non-survivors) (GEO: GSE167363)^33^. Similar to pediatric sepsis, cellular neighborhoods were significantly increased in neutrophil compartment in adult sepsis (**Fig. 7a**). Neutrophils in PBMCs are widely known as low-density neutrophils^34^. Conversely, fractions of other immune cells were decreased in abundance **(Fig. 7a** and **Suppl Fig. 10a)**. Neutrophils *RETN*, *CD24*, *CEACAM1*, and *CEACAM8* demonstrated similar expression profiles in both pediatric and adult sepsis **(Suppl Fig. 10b)**.

**Fig. 7:**
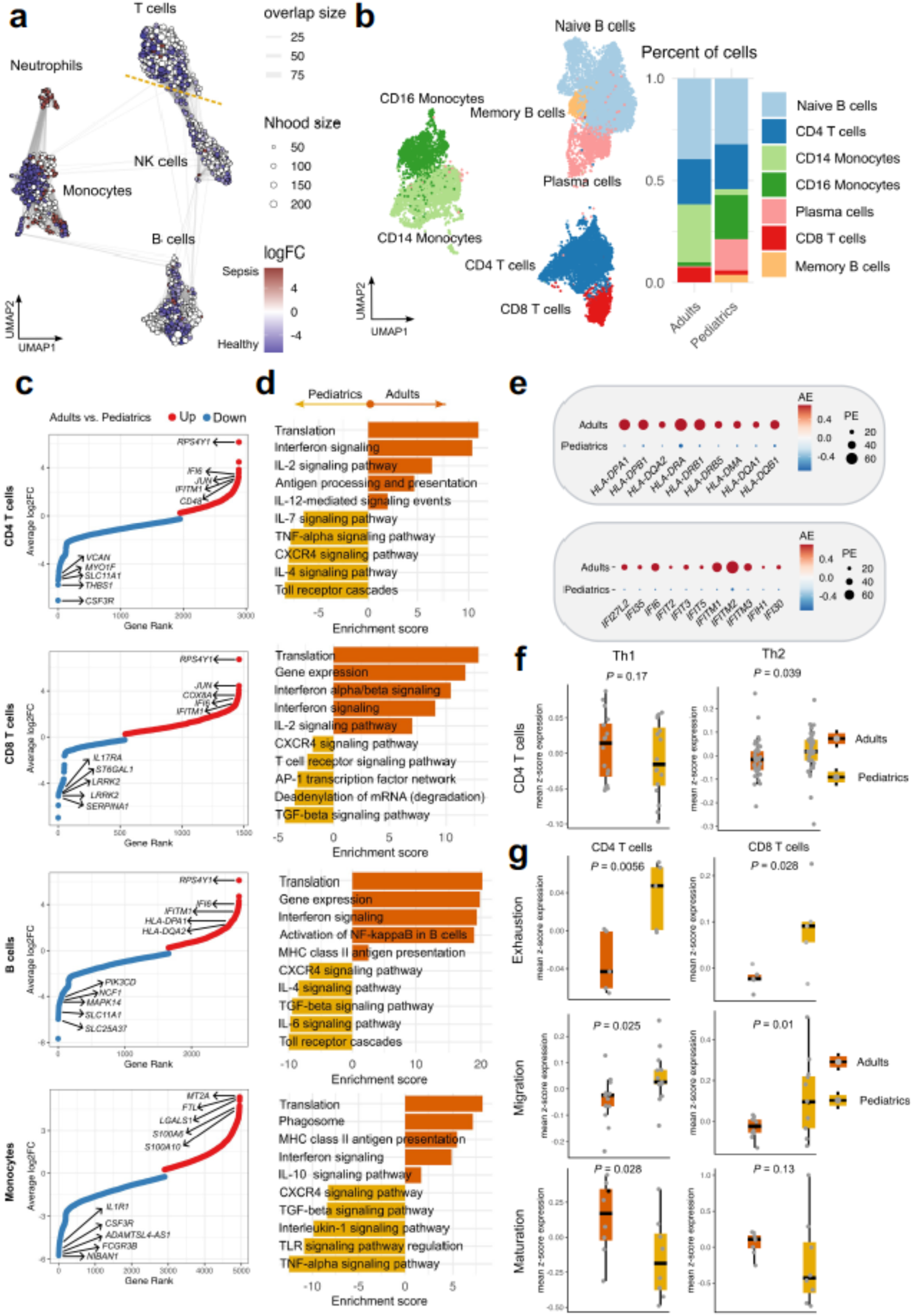
Comparison of major immune cells between Pediatric and adult sepsis. **a,** UMAP of differential cell abundance between adult sepsis (*n* = 5) and healthy donors (*n* = 2) with sampled neighborhoods colored by statistical significance (spatial FDR□<□0.05). **b,** UMAP for the major cell types of the merged acute phase samples of pediatrics (*n* = 6) and adults (*n* = 3) with their partitioning distribution. **c,** Gene rank plot for the DEGs (FDR□<□0.05) of the adults vs pediatrics pairwise comparison of CD4, CD8 T cells, B cells, and Monocytes. **d,** Bar plots of significant biological pathways associated with DEGs of the previous comparisons. **e,** Dot plots comparing expression levels of the interferon-stimulated genes (ISGs) and HLA-DR genes between adults and pediatric in total cells of sepsis acute phase. **f,** box plots comparing gene scores for Th1 and Th2 gene signatures between adults and pediatric in CD4 T cells of sepsis acute phase. Unpaired student t test was used for statistical analysis. **g,** Box plots comparing genes scores for exhaustion, migration, and maturation processes between adults and pediatric in CD4 T cells of sepsis acute phase. Unpaired student t test was used for statistical analysis.

We next integrated both single-cell datasets of adult (survivors) and pediatric sepsis in acute phase **(Fig. 7b)**. Although comparable fractions of CD4^+^ T cells were detected in both datasets, innate-like CD4 T cells deemed to be specific to pediatric sepsis **(Suppl Fig. 10c)**. We also observed a lower abundance of CD8^+^ T cells and enriched fractions of plasma and memory B cells in pediatric sepsis. Additionally, partitioning of CD14^+^ and CD16^+^ monocytes were more abundant in either adult or pediatric sepsis, respectively.

To gain a mechanistic understanding of the molecular differences between pediatric and adult sepsis, we carried out pairwise comparison of the major PBMC fractions **(Fig. 7c)**. Consequently, translation, interferon signaling, and antigen presenting pathways were significantly overrepresented in B, CD4^+^, CD8^+^ cells, and monocytes in adult sepsis **(Fig. 7d-e)**. Based on the enrichment of IL-4 signaling pathway in CD4^+^ T compartment in pediatric sepsis, CD4^+^ T cells represented more Th2 phenotype, but not Th1 compared to adult sepsis **(Fig. 7f).** We also observed higher exhaustion and migration scores in CD4 and CD8 cell fractions with lower maturity in pediatric sepsis **(Fig. 7g)**. These are consistent with the fact that pediatric T cells are still maturing and also prone to Th2 phenotype^6,25^. Whether or not these phenotypes represent escape routes for T cells to circumvent excessive inflammation during acute phase of sepsis in children needs to be examined in the future.

## Discussion

This study is considered the first work to decipher the complex cellular composition in pediatric sepsis at a single-cell level, charting alterations in frequencies and molecular states between acute and recovery phases. In our current work, pediatric sepsis resembled the well-recognized cellular phenomena in the periphery of adult sepsis including neutrophilia, lymphopenia, and dysregulation in antigen presentation^35^.

We have shown a discrete shift in the functional neutrophil plasticity from a suppressive state to mature and effector states^36^, facilitated by a global switch in the cellular composition of neutrophils rather than by a transcription reprogramming. This could be attributed to their fleeting lifespan and reduced transcription activity^37,38^. We were also able to detect *RETN* gene as a regulatory core that plays a crucial role in sepsis pathogenesis^39^. Resistin was found to mediate detrimental effects by impairing neutrophil migration, ROS production, and intracellular bacterial clearance in a dose-dependent manner^40,41^. In our analysis, the *RETN*-expressing neutrophil subpopulation was characterized by overexpression of genes associated with surface proteins such as CD24, CD66a, and CD66b. A recent study by J. Kwok et al reported that CD66b^+^ sepsis neutrophils caused suppression of T cells^35^, which could be directly linked to CD66a (*CEACAM1*), a heterophilic ligand for the T cell inhibitory molecule T-cell immunoglobulin domain and mucin domain-3 (*TIM3*)^42^. This is also in line with our data in both human and murine sepsis, in which the biological processes enriched in the *CEACAM1* gene co-expression network were assigned to negative regulation of T cell activation. In addition, the strong correlation between expression levels of *RETN* and *CEACAM1* in the acute sepsis phase could reflect the deleterious clinical outcomes in this phase^43^.

The appearance of innate-like CD4 T cells in our current work was first reported by Gibbons et al as *CXC8L*-expressing CD4 T cells in newborn infants that provided a defense mechanism against infections^29^. This is in concert with our data that these cells solely sustained in memory CD4 T compartment in acute sepsis phase. Additionally, since CD31/*PECAM1* marker was weakly expressed by innate-like CD4 T cells, we were able to define them as CSF3R^+^ T cells. One possible explanation for their presence is the defect in stringent thymic selection of CD4 T cells under stress in pediatric subject. This was evidenced by our data through the lower abundance and absence of innate-like CD4 T cells in older children (school age) and adult subjects, respectively. It is worth noting that *CXCL8* (IL-8) has been widely used as prognostic marker for mortality in the Pediatric Sepsis Biomarker Risk Model (PERSEVERE)^44,45^.

Clearly T cell comparison between adult and pediatric sepsis showed distinctive differences in adaptive immune cells. There is an unmet need to conduct further research in the future to explore the biological and functional roles of these cells.

The utility of (monocyte) mHLA-DR expression in the periphery has been well established in diagnosis of adult sepsis with a threshold < 30 % to be linked with increased mortality^46^, meanwhile similar threshold might not be applicable in pediatric sepsis^47^. Our data illustrated an enhanced HLA-DR expression in adult sepsis compared to pediatric sepsis, indicating that such a threshold could be even lower in pediatrics. In parallel, the overexpression of interferon-stimulated genes (ISGs) in adult sepsis was in line with several studies, in which stimulating monocytes with IFN improved HLA-DR expression^48–51^. Our study serves as a rich source of information about the phenotypic diversity of immune cells in the periphery of pediatric sepsis individuals, which could be exploited by future translation studies.

## Acknowledgements

We thank the patients and healthy donors and their parents for the participation in this study. We also thank Samuel Kim, Sheng Xiang Huang, Karina Lukovits, and Rachel Bernier (all Boston Children’s Hospital) for the technical assistance.

## Contributions

F.A, S.K. and K.Y. developed the research concept. S.K. and K.Y. provide the funding. S.K. and K.Y. carried out the *in vivo*/*ex vivo* human and mice experiments. K.Y. curated the raw single-cell data. F.A. performed bioinformatic analysis for bulk and single-cell RNA-Seq data. F.A. conducted *in vitro* experiments. F.A. and K.Y wrote and edited the original manuscript draft.

