## Supplementary material for "The landscape of immune dysregulation in pediatric sepsis at a single-cell resolution": Supple Figures

**a**

| Sample | Age (year) | Gender | pSOFA |
|---|---|---|---|
| AP1 | 1 | F | 6 |
| AP2 | 0.7 | M | 3 |
| AP3 | 15 | F | 12 |
| AP4 | 15 | M | 2 |
| AP5 | 3 | F | 7 |
| AP6 | 15 | M | 8 |
| RP1 | 1 | F | 2 |
| RP2 | 0.7 | M | 0 |
| RP3 | 15 | F | 4 |
| RP4 | 15 | M | 0 |
| RP5 | 3 | F | 0 |
| RP6 | 15 | M | 3 |
| HC1 | 0.3 | M | 0 |
| HC2 | 10 | F | 0 |
| HC3 | 6 | M | 0 |
| HC4 | 0.8 | F | 0 |

**b**

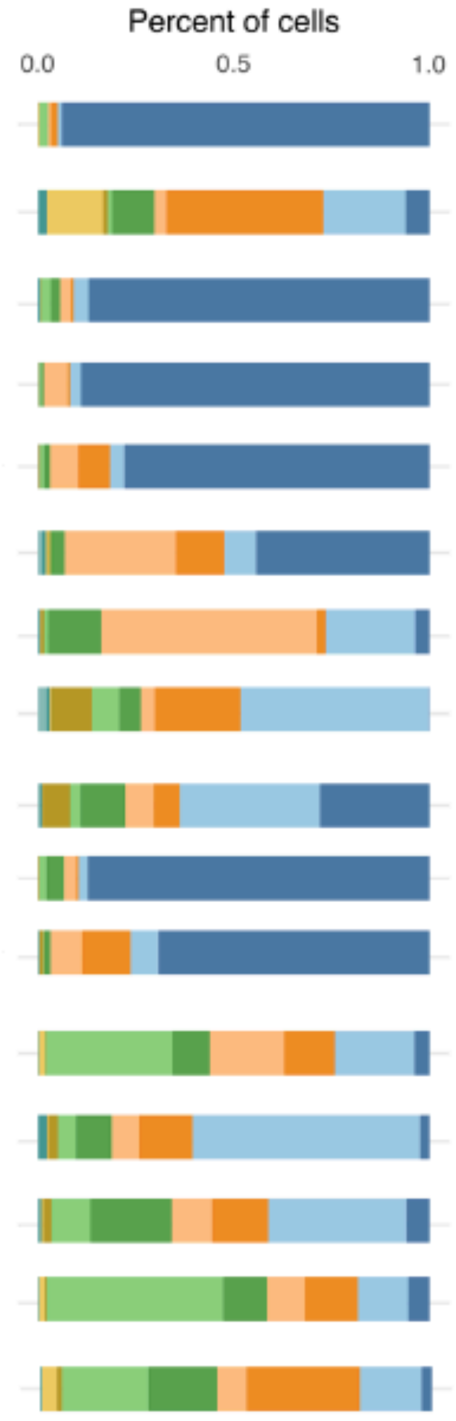


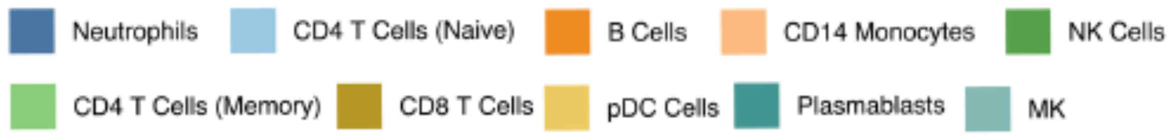


**Supplementary Fig. 1: Demographic characteristics of study subjects. a**, Age, gender, and pSOFA scores of the study subjects. **b**, Partitioning distribution of major cell types for each donor of the study subjects.

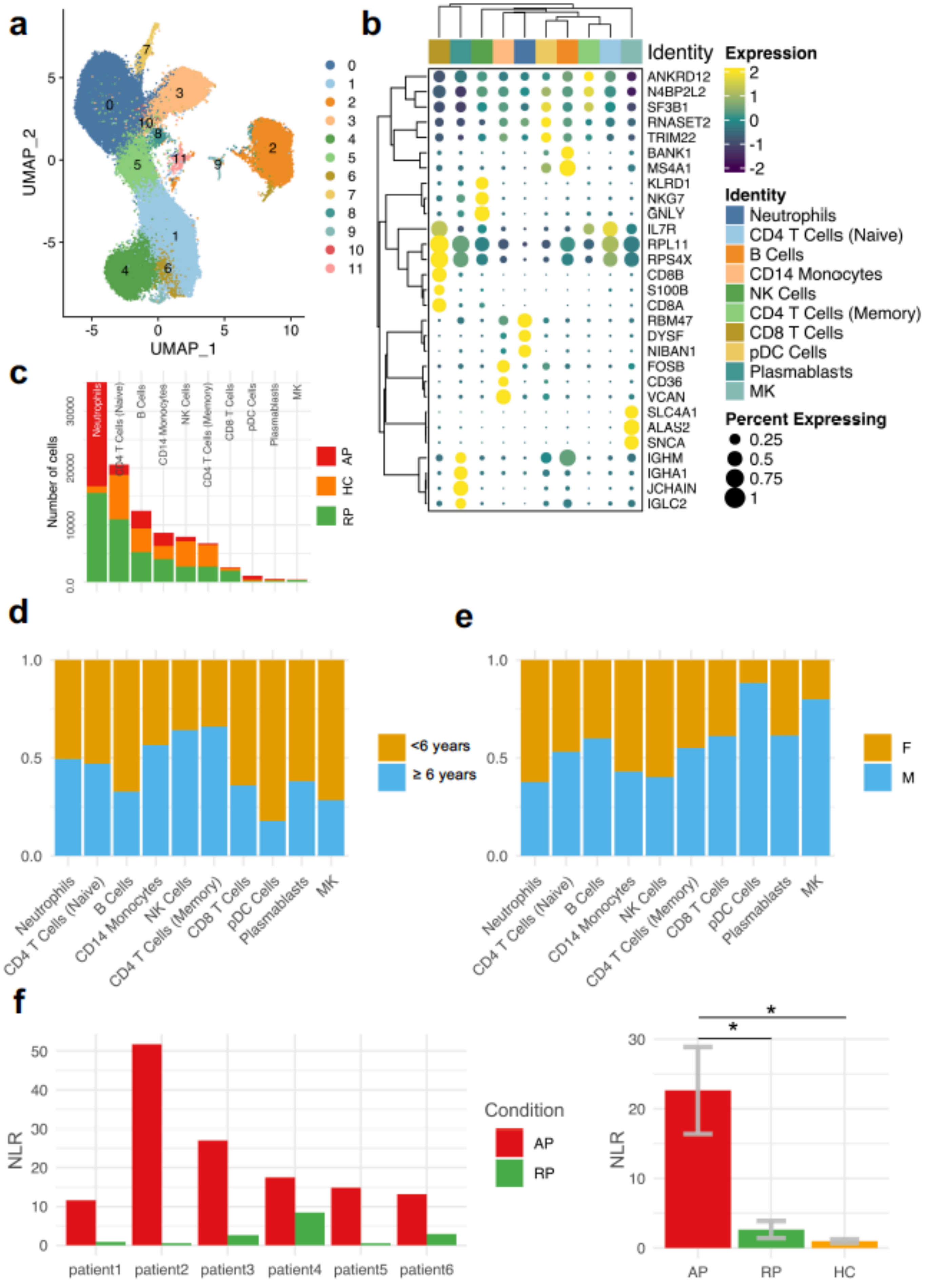

**Supplementary Fig. 2: Seurat clustering and NLR scores comparison. a,** Original Seurat clustering at a 0.3 resolution factor. **b,** Clustered heatmap of the top three genes of each major cell type. **c,** Partitioning distribution of the major cell types across study subject conditions as absolute counts. **d,** Partitioning distribution of the major cell types across gender groups (M = Male; F = Female). **e,** Partitioning distribution of the major cell types across age groups (< 6 years and ≥ 6 years). **f,** Comparison of NLR scores for each pediatric sepsis patient in AP and RP (left) and combined comparison of NLR scores across the three study subject conditions (right). One-way ANOVA with Bonferroni *post hoc* analysis was performed for statistical analysis. * p< 0.05.

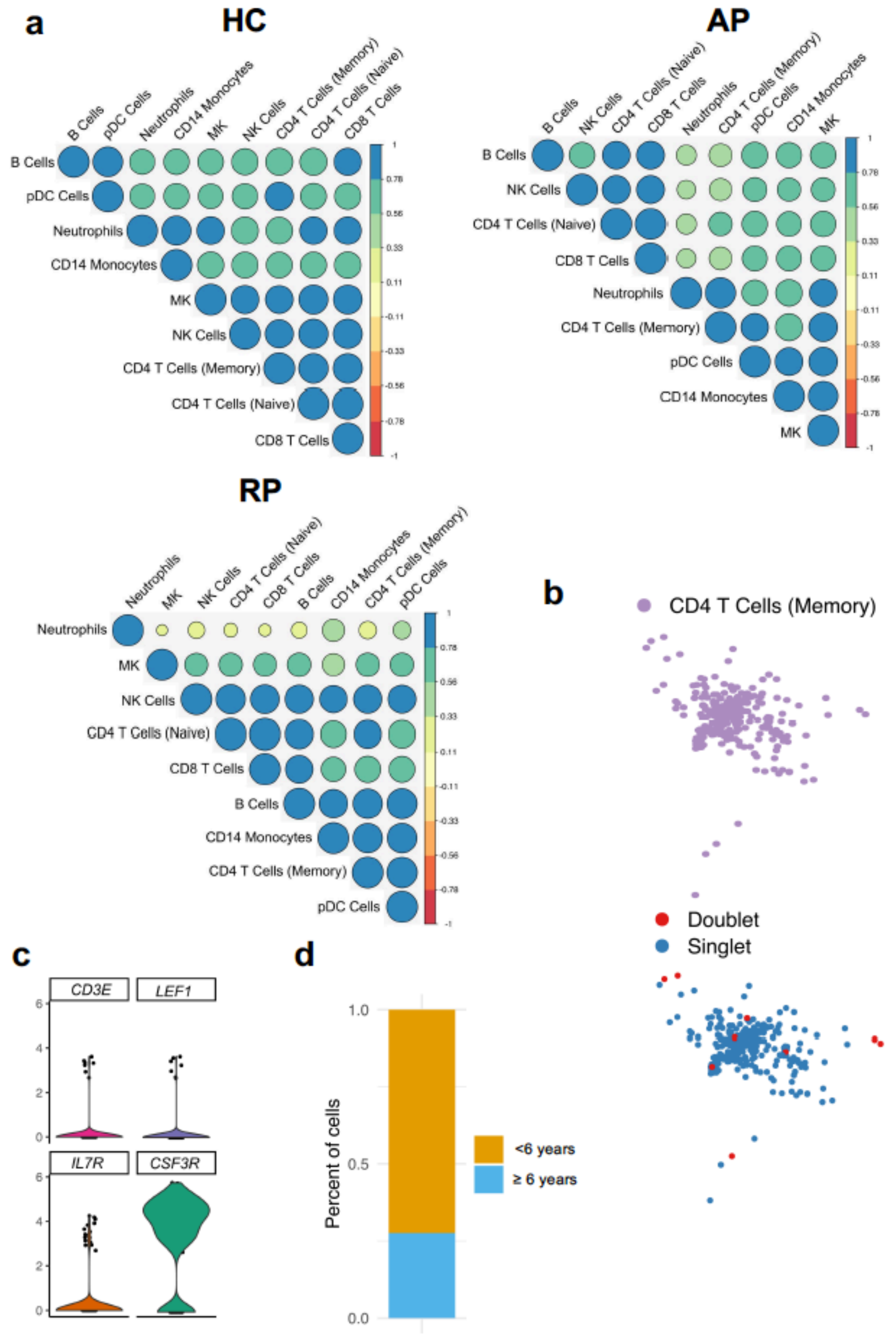


**Supplementary Fig. 3: Characterization of innate-like CD4 T cells. a,** Hierarchal clustering analysis of the top 500 variable genes of the major cell types across the three study subject conditions. **b,** Cell-doublet analysis of CD4 T cells (memory) compartment using doublet finder algorithm with default sittings. **c,** Violin plot of the basic CD4 T cell marker genes as well as CSF3R as marker gene in innate-like CD4 T cells (previously known as memory CD4 T cells). **d,** Partitioning distribution of the major cell types across age groups.

**a**

| N1 | N2 | N3 | N4 | N5 | N6 | N7 | N8 | N9 |
|---|---|---|---|---|---|---|---|---|
| *HNRNPH1* | *CSF3R* | *PI3* | *MMP9* | *ADAMTSL4-AS1* | *MME* | *RBP7* | *ZDHHC19* | *LCN2* |
| *TSC22D3* | *AC007271.1* | *EPHB1* | *NCF1* | *CAMK1D* | *FCGR3B* | *PPP1R12B* | *DACH1* | *LTF* |
| *MICAL1* | *CD177* | *PPCDC* | *PADI4* | *AP001977.1* | *CXCR2* | *FCGR3B* | *TRPS1* | *CAMP* |
| *PGGHG* | *IL1R1* | *AC022217.3* | *S100A12* | *SCLT1* | *S100A11* | *CDA* | *CD177* | *ABCA13* |
| *KIAA1551* | *KCNMA1* | *MICAL2* | *TYMP* | *LUCAT1* | *IFITM2* | *FTL* | *ARG1* | *CD24* |
| *DUSP1* | *SLC25A37* | *SCLT1* | *ATP6V0C* | *AC005050.3* | *G0S2* | *LAMTOR4* | *ST6GALNAC3* | *MMP8* |
| *FOSB* | *ZDHHC19* | *WLS* | *TSPO* | *SSH2* | *FTH1* | *FCN1* | *FGD4* | *CEACAM8* |
| *LENG8* | *JAK3* | *AP002381.2* | *JAK3* | *EPHB1* | *S100A4* | *GRN* | *DYSF* | *BPI* |
| *FOS* | *BCL6* | *MAML2* | *PLBD1* | *AC099489.1* | *B2M* | *HMGB2* | *MAPK14* | *RETN* |
| *SAT1* | *MAPK14* | *LUCAT1* | *NTNG2* | *MAML2* | *FAM129A* | *LYZ* | *SYNE1* | *STOM* |

**b**

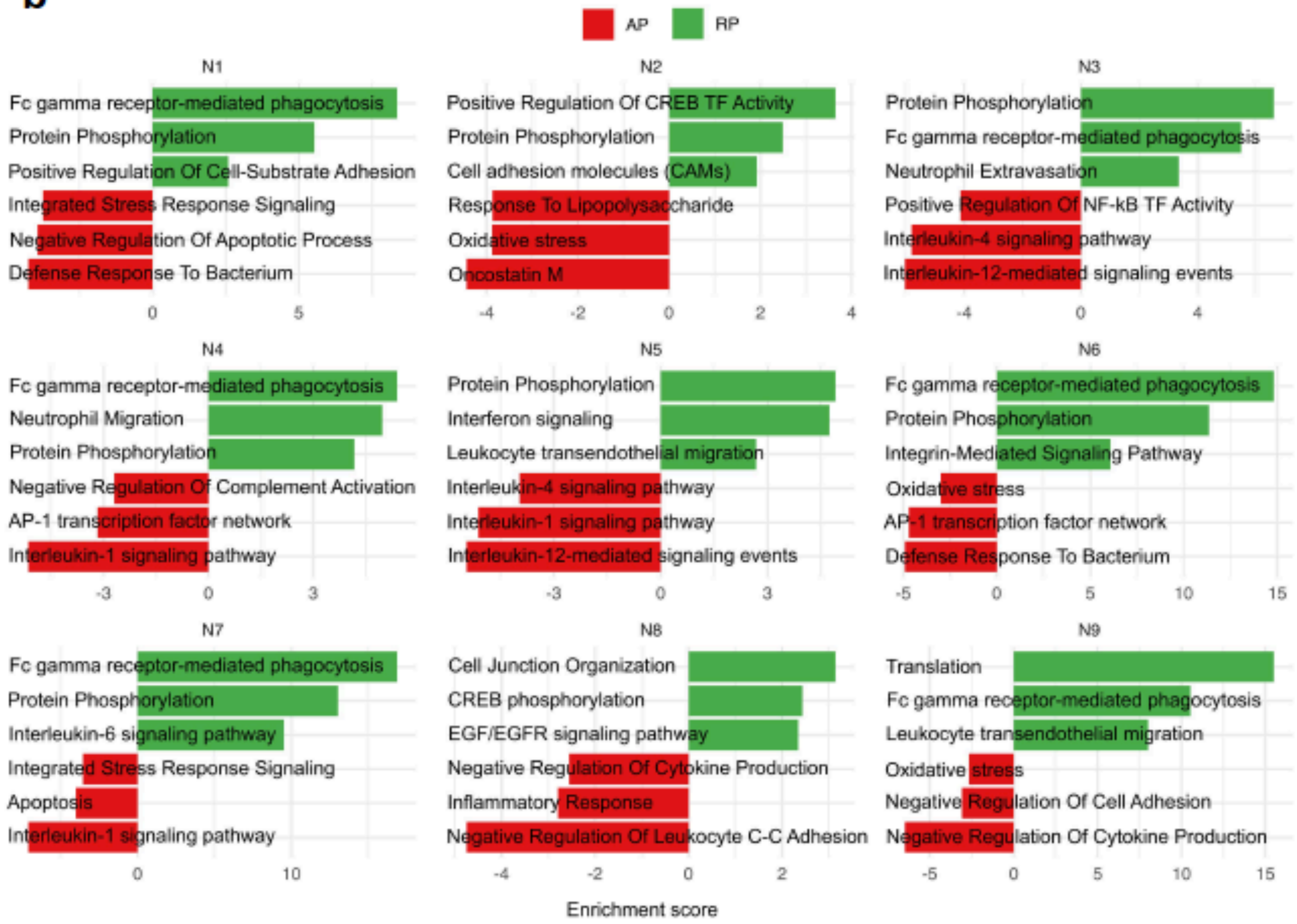


**Supplementary Fig. 4: Top genes, biological processes, and pathways of neutrophil subpopulations. a,** Top 10 genes for each of the nine neutrophil subpopulations. **b,** Significant biological pathways and processes for each neutrophil subcluster utilizing the DEGs between AP and RP.

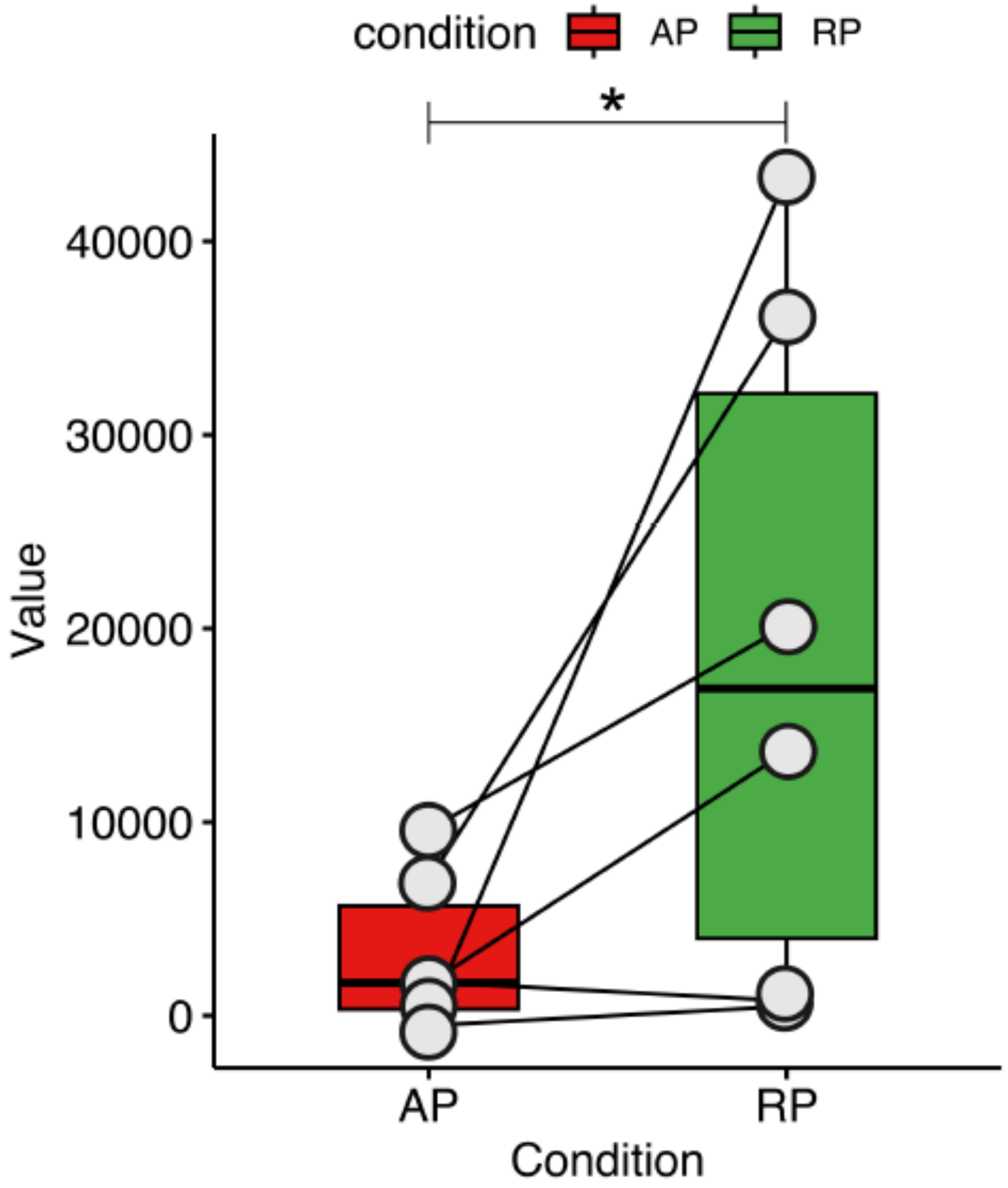


**Supplementary Fig. 5: Phagocytosis scores for the two sepsis phases.** Box plot showing the paired comparison of the phagocytosis scores between AP and RP. Paired t test comparing phagocytosis scores. *$P < 0.05$.

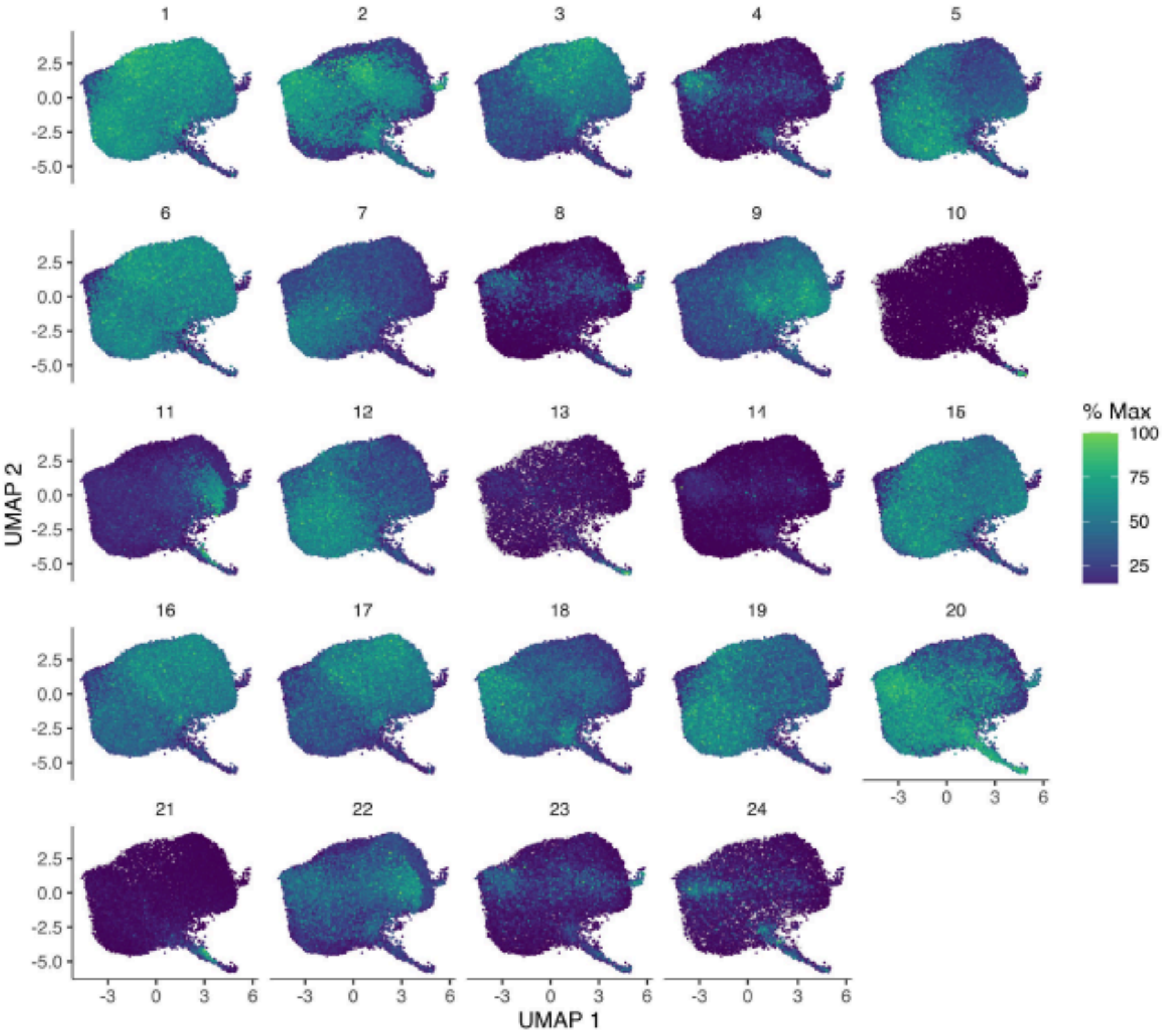


**Supplementary Fig. 6: Gene modules of the neutrophils trajectory.** 24 gene modules across total neutrophil population computed with Monocle 3 algorithm at a resolution = $10^{-3}$.

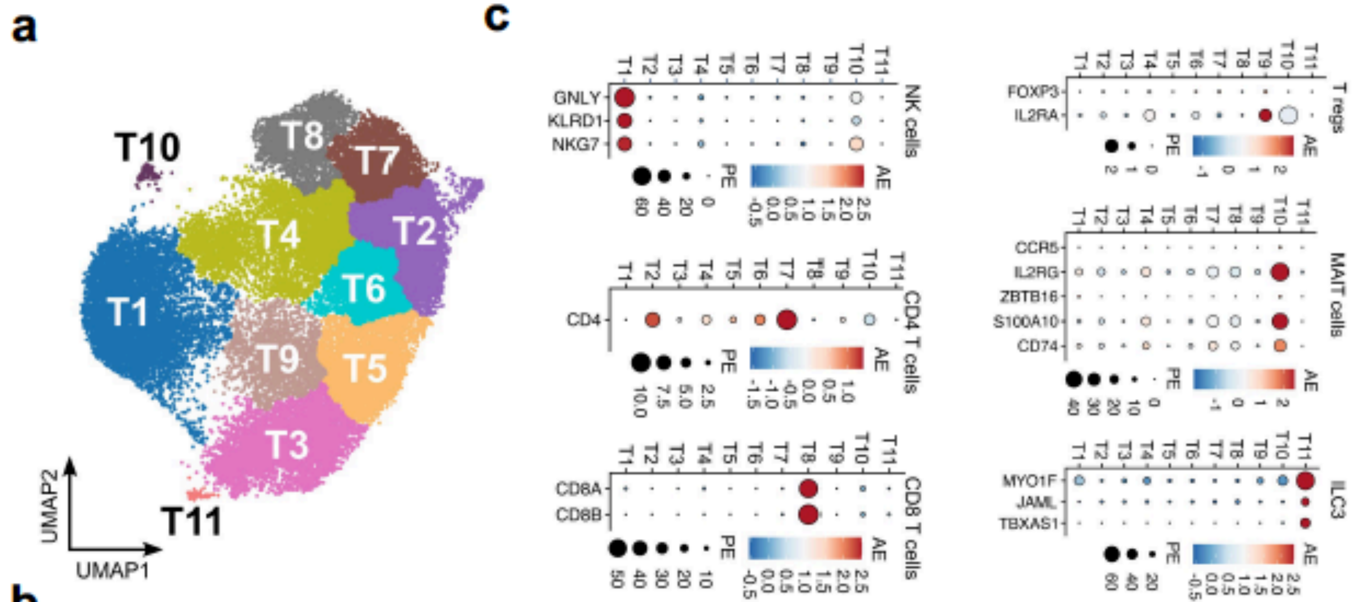


**b**

| T1 | T2 | T3 | T4 | T5 | T6 | T7 | T8 | T9 | T10 | T11 |
|---|---|---|---|---|---|---|---|---|---|---|
| *GNLY* | *CCR7* | *RIPOR2* | *S100A4* | *LEF1* | *LTB* | *NACA* | *CD8B* | *RORA* | *TOP2A* | *DYSF* |
| *SPON2* | *TPT1* | *PLCG2* | *AHNAK* | *CAMK4* | *TCF7* | *BTF3* | *CD8A* | *SYNE2* | *MKI67* | *NIBAN1* |
| *KLRD1* | *FAU* | *S100A9* | *IL32* | *IL6ST* | *FAU* | *LDHB* | *S100B* | *NKTR* | *CENPF* | *RBM47* |
| *NKG7* | *TMSB10* | *NAMPT* | *CD52* | *BCL11B* | *CAMK4* | *CD3D* | *NELL2* | *MACF1* | *TYMS* | *GLT1D1* |
| *PRF1* | *EEF1B2* | *DDX17* | *S100A6* | *HNRNPH1* | *FOXP1* | *TMSB10* | *NUCB2* | *DDX17* | *RRM2* | *GAB2* |
| *ZEB2* | *GIMAP7* | *DDX5* | *RORA* | *DDX17* | *LEF1* | *GIMAP7* | *NACA* | *CXCR4* | *NUSAP1* | *WDFY3* |
| *CCL5* | *LDHB* | *SMCHD1* | *LTB* | *IL7R* | *NOP53* | *EIF3K* | *CD3D* | *SAT1* | *STMN1* | *MAML3* |
| *EFHD2* | *LEF1* | *USP15* | *EEF1A1* | *FYB1* | *SARAF* | *COX7C* | *NPM1* | *S100A9* | *TUBA1B* | *ACSL1* |
| *MYO1F* | *LTB* | *S100A8* | *TMSB4X* | *ATM* | *EEF2* | *RACK1* | *COX4I1* | *CCNL1* | *TUBB* | *RNF24* |
| *RUNX3* | *NACA* | *SORL1* | *B2M* | *RIPOR2* | *EIF4B* | *COMMD6* | *CD7* | *CELF2* | *PCNA* | *LYN* |

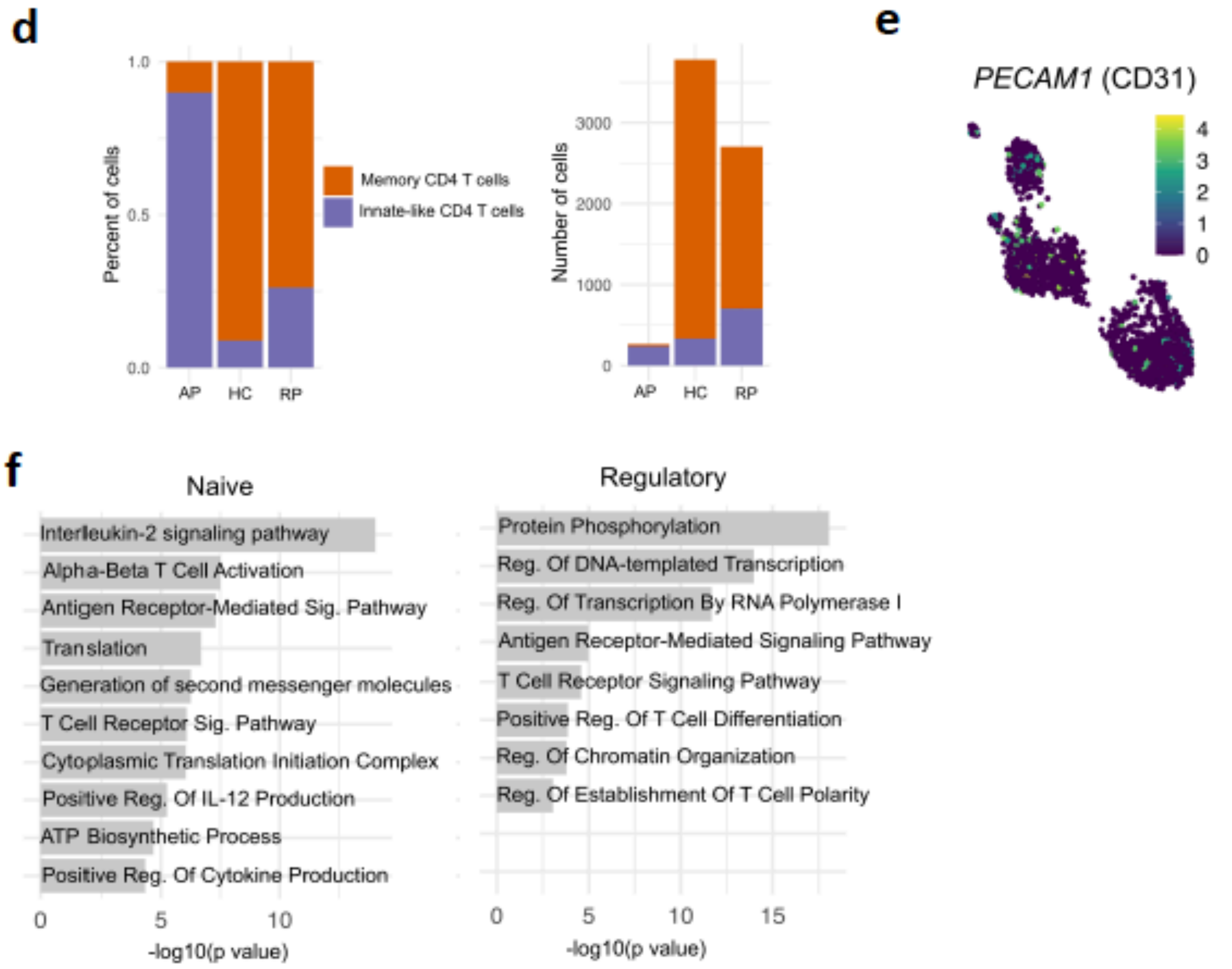


**Supplementary Fig. 7: Top genes, biological processes, and pathways of T and NK subpopulations. a,** UMAP showing 11 subclusters of T and NK subclusters. **b,** Top 10 genes for each of the 11 T and NK subpopulations. **c,** Dot plots of the gene markers annotating the subpopulations of T and NK cells. **d,** Partitioning distribution of the memory and innate-like CD4 T cells across the three study subject conditions. **e,** Feature plot for the expression of *PECAM1* (CD31) as traditional marker gene for innate-like (*CXCL8*-expressing) CD4 T cells. **f,** Bar plots showing the significant biological processes and pathways associated with the significant gene markers for naïve and regulatory CD4 T cells in AP (FDR < 0.05).

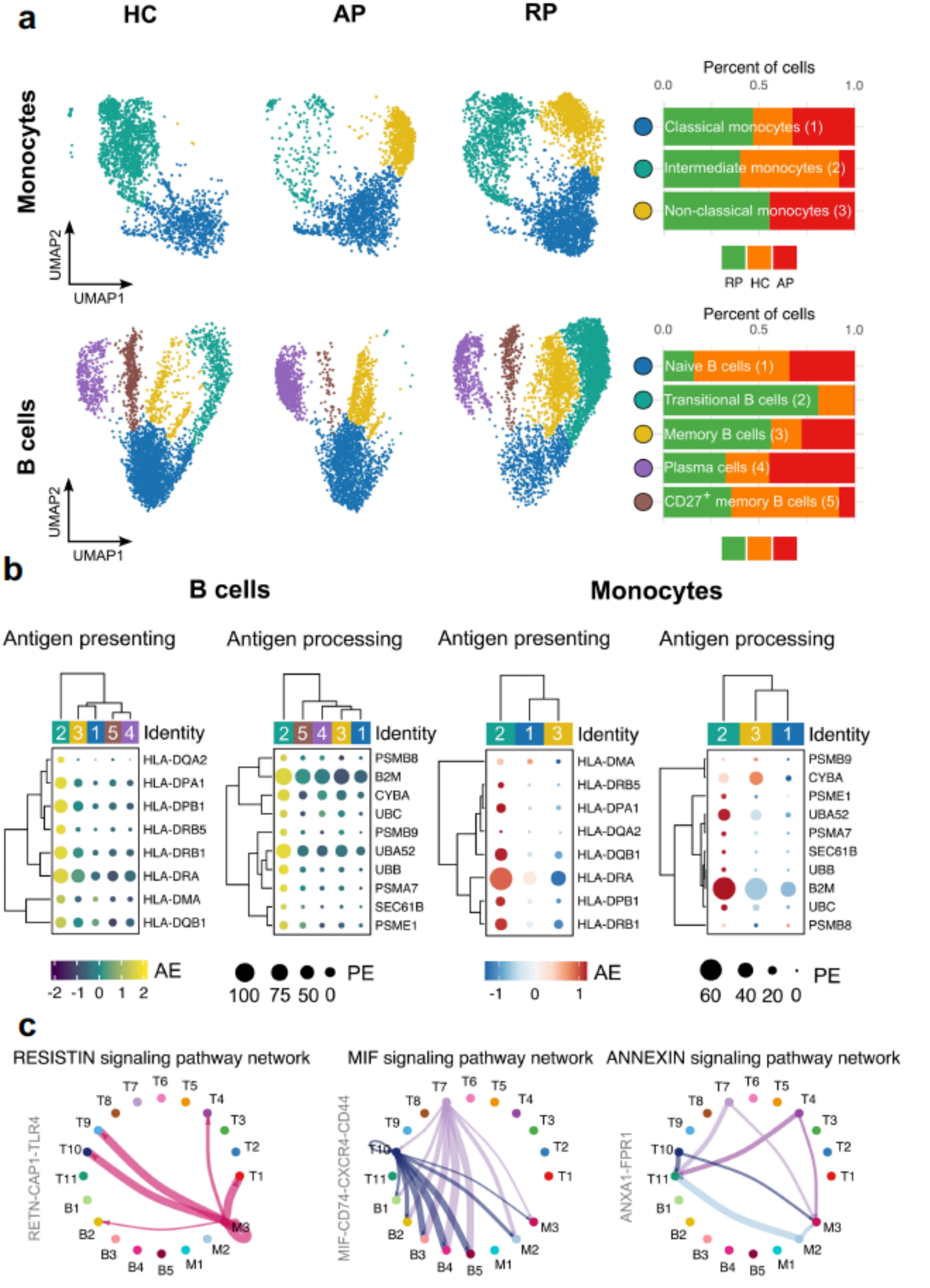


**Supplementary Fig. 8: Lack of antigen presenting and processing genes in monocytes and B cells of AP. a,** UMAPs and bar plots depicting the abundance differences in monocytes (up) and B cells (down) across the three study subject conditions. **b,** Clustered dot plots showing the expression of the antigen presenting and processing segregated across the subpopulations of B cells (right) and monocytes (left). **c,** Cell-cell communication analysis between subpopulations of T, B cells, and monocytes showing the RESISTIN signaling interaction enriched in AP (left) and MIF and ANNEXIN signaling enriched in RP (right).

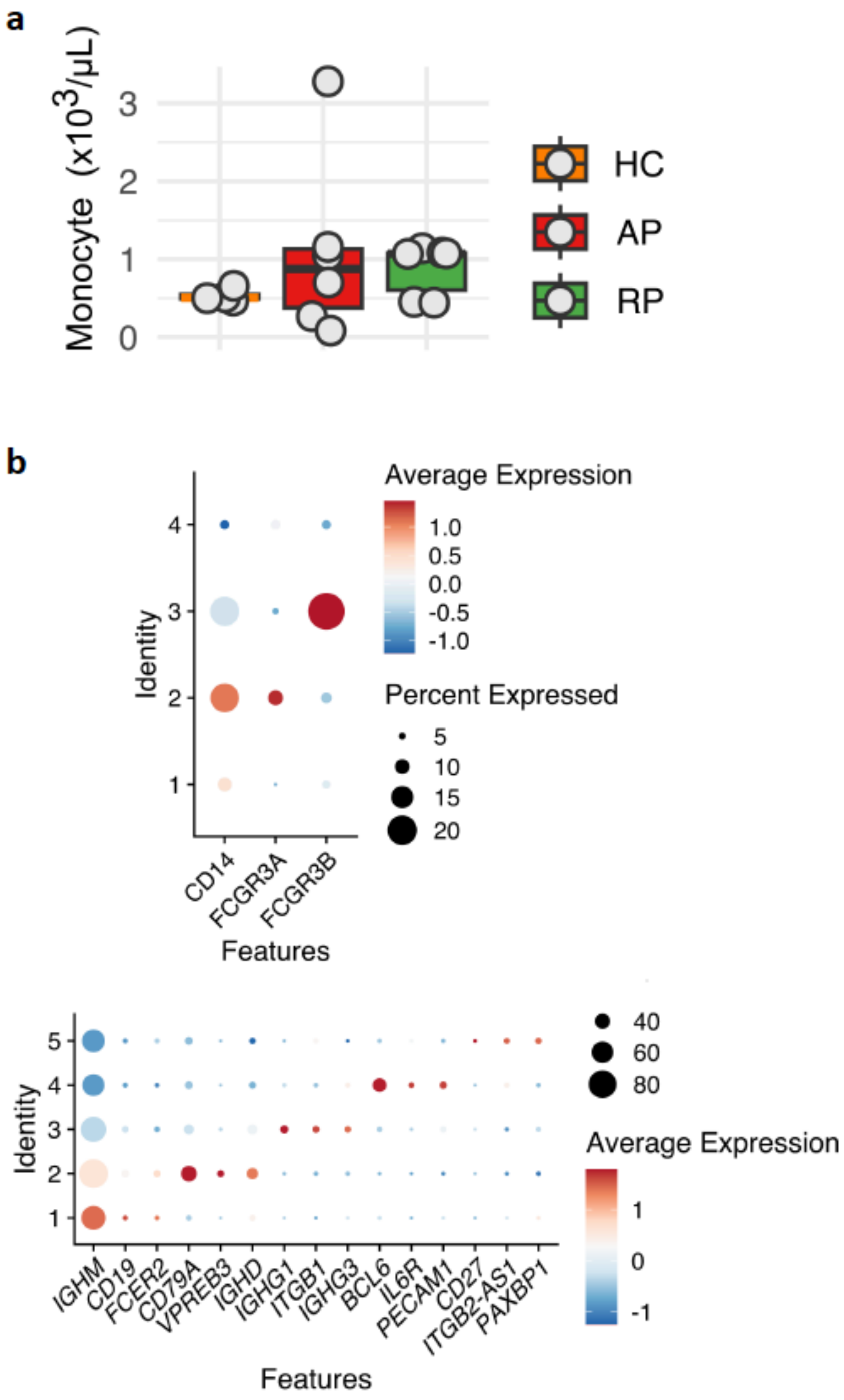


**Supplementary Fig. 9: Annotation of monocytes and B cells subpopulations. a,** Box plot showing the comparison of the monocyte counts between the three study conditions. One-way ANOVA with Bonferroni *post hoc* analysis was performed for statistical analysis. No statistical significance was noted. **b,** Dot plots of the gene markers annotating the subpopulations of monocytes (up) and B cells (down).

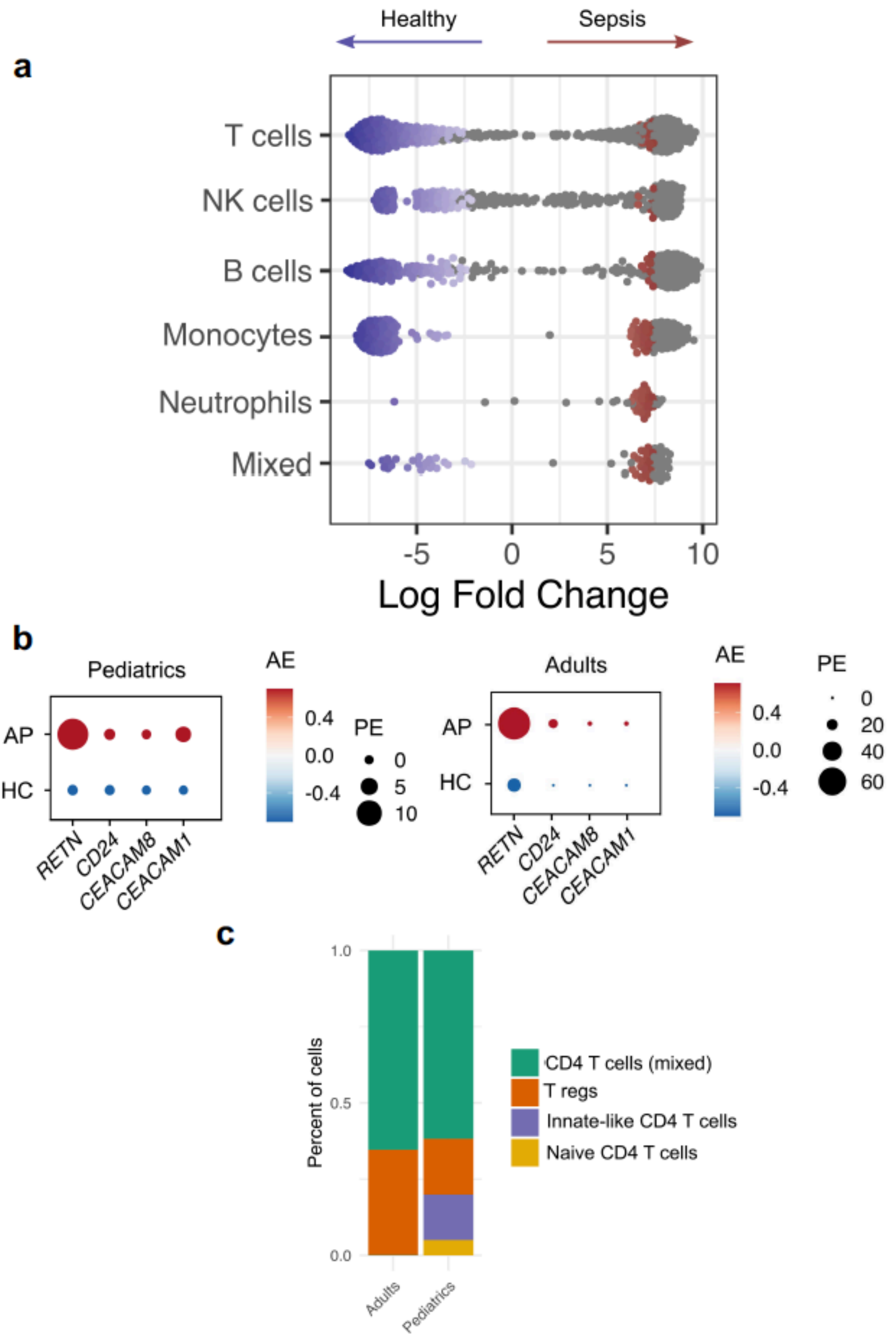


**Supplementary Fig. 10: Comparison of neutrophil gene signatures between pediatric and adult sepsis acute phase. a,** Beeswarm plot of differential cell abundance in major cell types of PBMCs between sepsis ($n$ = 5) and healthy ($n$ = 2) adult individuals. **b,** Dot plots comparing four neutrophil signature genes between AP and HC in pediatric and adult individuals. **c,** Partitioning distribution of CD4 T cell subpopulations across pediatric and adult individuals.
